# A heterodimeric SNX4:SNX7 SNX-BAR autophagy complex coordinates ATG9A trafficking for efficient autophagosome assembly

**DOI:** 10.1101/2020.03.15.990713

**Authors:** Zuriñe Antón, Virginie M.S. Betin, Boris Simonetti, Colin J. Traer, Naomi Attar, Peter J. Cullen, Jon D. Lane

## Abstract

Efficient mammalian autophagosome biogenesis requires coordinated input from other cellular endomembrane compartments. Such coordination includes the stimulated trafficking to autophagosome assembly sites of the essential autophagy proteins, ATG9 and ATG16L1, via distinct endosomal compartments. Protein trafficking within the endocytic network is directed by a conserved family of sorting nexins (SNXs), with previous studies implicating SNX18 (an SH3 domain-type SNX-BAR protein) in the mobilisation of ATG9A and ATG16L1 from recycling endosomes during autophagy. Using siRNA and CRISPR-Cas9, we demonstrate that a second mammalian SNX-BAR, SNX4, is needed for efficient LC3 lipidation and autophagosome assembly in mammalian cells. SNX-BARs exist as homo- and heterodimers, and we show that SNX4 forms functional heterodimers with either SNX7 or SNX30, and that these associate with tubulovesicular endocytic membranes at steady state. Detailed image-based analysis during the early stages of autophagosome assembly reveal that SNX4:SNX7 is the autophagy-specific heterodimeric SNX-BAR complex, required for efficient recruitment/retention of core autophagy regulators at the nascent isolation membrane. SNX4 partially co-localises with juxtanuclear ATG9A-positive membranes, with our data linking the SNX4 autophagy defect to the mis-trafficking and/or retention of ATG9A in the Golgi region. Together, our findings show that the SNX4:SNX7 heterodimer coordinates ATG9A trafficking within the endocytic network to establish productive autophagosome assembly sites.

**SUMMARY STATEMENT:** A heterodimeric SNX4:SNX7 SNX-BAR complex regulates mammalian autophagosome assembly through the control of ATG9 trafficking.

## INTRODUCTION

Macroautophagy (herein referred to as “autophagy”) describes the sequestration of cytoplasmic material within double membrane-bound vesicles (autophagosomes) that deliver their contents to lysosomes for degradation and recycling. Autophagosome assembly requires the concerted actions of conserved ATG proteins that are targeted to specialised endoplasmic reticulum (ER) sub-domains (called omegasomes) that are enriched in the phosphoinositide, PtdIns(3)P (Axe et al., 2008). Consequently, molecules with affinity for either PtdIns(3)P and/or curved membrane profiles (e.g. Bin/Amphiphysin/Rvs [BAR] domain-containing proteins (McMahon and Boucrot, 2015)) have been implicated in the control of autophagosome biogenesis. The WIPI family of PtdIns(3)P effector proteins are essential mediators of autophagosome assembly, coupling localised PtdIns(3)P to the recruitment of the ATG8 (LC3/GABARAP) lipidation machinery via direct binding (in the case of WIPI2b) to ATG16L1 (Dooley et al., 2015). Also acting during the autophagosome expansion phase is the BAR domain-containing protein, SH3GLB1 (BIF-1/Endophilin B1), which binds to UVRAG/Beclin-1 to stimulate the autophagy PIK3C3/VPS34 kinase, whilst also facilitating trafficking of ATG9 to the autophagosome assembly site (Takahashi et al., 2007; Takahashi et al., 2011). Identification of further PtdIns(3)P effectors and/or BAR domain-containing proteins with the potential to influence autophagosome expansion/shaping remains a key objective.

Sorting nexins (SNXs) are a family of peripheral membrane proteins defined by the presence of a PX (phox homology) domain (Carlton et al., 2004; Cullen, 2008; Seet and Hong, 2006; Teasdale and Collins, 2012; Teasdale et al., 2001), and of the 33 SNXs annotated in the human genome, many interact with PtdIns(3)P. As this lipid is enriched on early elements of the endocytic network (Gillooly et al., 2000), most SNXs are targeted to the cytosolic face of membrane bound compartments that make up this diverse organelle. For one evolutionary conserved sub-family of SNXs—the SNX-BARs (Carlton et al., 2004; Habermann, 2004)—the presence of an additional carboxy-terminal BAR domain conveys upon them the ability to generate and/or stabilize membrane tubules (Carlton et al., 2004). Mammalian cells possess twelve SNX-BAR family members—SNX1, SNX2, SNX4 through to SNX9, SNX18, SNX30, SNX32 and SNX33 (van Weering and Cullen, 2014; van Weering et al., 2010)—within which there is emerging evidence for a restricted pattern of BAR domain-mediated homo- and hetero-dimerisations (van Weering and Cullen, 2014; van Weering et al., 2010). Thus the SH3 domain-containing SNX-BARs—SNX9, SNX18 and SNX33—homodimerise to coordinate actin polymerization with vesicle scission at sites of high membrane curvature (Dislich et al., 2011; Haberg et al., 2008; Lundmark and Carlsson, 2009; Schoneberg et al., 2017; van Weering et al., 2012) (although this conclusion remains controversial (Park et al., 2010)). In contrast, SNX1, SNX2, SNX5, and SNX6 (and its neuronal counterpart SNX32) make up a membrane re-sculpturing coat complex, named ESCPE-1 that consists of heterodimers of SNX1:SNX5, SNX1:SNX6, SNX2:SNX5 and SNX2:SNX6 (Simonetti et al., 2017; Simonetti et al., 2019).

Genetic screens in yeast have implicated the SNX-BARs, *SNX4* (also known as *ATG24B* and *CVT13*) and *SNX42* (also known as *ATG20* and *CVT20*), as regulators of selective autophagy, including cytoplasm-to-vacuole targeting (cvt) (Nice et al., 2002) (reviewed in (Lynch-Day and Klionsky, 2010)), pexophagy (Ano et al., 2005; Deng et al., 2012), mitophagy (Kanki et al., 2009; Mendl et al., 2011), and selective degradation of fatty acid synthase (Shpilka et al., 2015). Snx4 co-localises with Atg8 at the PAS (Zhao et al., 2016), and perturbing the PtdIns(3)P-binding capabilities of these proteins prevents their association with the phagophore assembly site (PAS), and impairs the Cvt pathway (Nice et al., 2002). Snx4 interacts with Atg17 (the yeast equivalent of mammalian RB1CC1/FIP200) (Nice et al., 2002; Uetz et al., 2000), a protein that regulates Atg1-stimulated Atg9 trafficking to the PAS (Sekito et al., 2009). Taking this further, Popelka and colleagues argued that yeast Atg11 assembles with Snx4/Atg24 and Atg20, replacing the non-selective autophagy Atg1 sub-complex (Atg17-Atg31-Atg29) to mediate selective autophagy (Popelka et al., 2017). Atg11 is a scaffolding protein that is specific for selective forms of autophagy in yeast (Zientara-Rytter and Subramani, 2019). In other hands, defects in non-selective autophagy have been reported in Snx4-null yeast (Ma et al., 2018), meanwhile deletion of *ATG24B or ATG20* in the background of other Golgi/endosomal mutants resulted in synthetic starvation-induced (non-selective) autophagy defects, suggesting compensatory masking of phenotypes in single deletion settings (Ohashi and Munro, 2010). In yeast, a series of dimeric interactions defined by weak Snx4:Snx4 homodimers and more pronounced Snx4:Snx41 and Snx4:Snx42 heterodimers have been described (Hettema et al., 2003) (see also (Ito et al., 2001; Popelka et al., 2017; Uetz et al., 2000; Vollert and Uetz, 2004)), and these findings are consistent with data obtained using recombinant human proteins (Traer et al., 2007). Which of mammalian SNX7 and SNX30 is the functional homologue of yeast Snx41 and Snx42 is difficult to establish given their respective sequence similarities, and precise roles for homo- or heterodimeric complexes established within this group of proteins remain uncertain. Phylogeny and dimerization patterns suggest that Snx42/Atg20 is likely to be the yeast equivalent of mammalian SNX30 (Popelka et al., 2017), and intriguingly, an indirect role for Snx4:Snx42 during autophagosome-to-vacuolar fusion via coordinated mobilisation of phosphatidylserine-containing membranes from the endocytic compartment has been described (Ma et al., 2018).

An imaging-based LC3 lipidation screen has described a role for the SH3-containing SNX-BAR, SNX18, during autophagy in mammalian cells (Knaevelsrud et al., 2013). SNX18 contains a conserved LIR (LC3-interacting) motif, and binds dynamin-2 independently of the LIR to mediate ATG9A trafficking from the recycling endosome and ATG16L1- and LC3-positive membrane delivery to the autophagosome assembly site (Knaevelsrud et al., 2013; Soreng et al., 2018). Here, we have tested whether further human SNXs contribute during autophagy, and we identify SNX4 as a positive autophagy regulator. Further, we have investigated the concept of restricted patterns of dimeric interactions within the mammalian SNX-BAR family, asking how this behaviour modulates the autophagy response with respect to SNX4. We present data establishing SNX4 as a core component of two heterodimeric endosomal associated complexes described by SNX4:SNX7 and SNX4:SNX30. Moreover, we show that the SNX4:SNX7 heterodimer is a positive regulator of autophagosome assembly in mammalian cells. Our data suggest that SNX4 complexes promote autophagosome assembly kinetics by mobilising ATG9A-associated membranes from the juxtanuclear area of the cell in response to autophagy stimulus.

## RESULTS

### siRNA suppression of SNX4 expression impairs autophagy

Given the evidence implicating Snx4 in various forms of autophagy in yeast, we tested for possible roles for mammalian SNX4 during amino acid/growth factor starvation-induced autophagy in cell culture. Immunoblotting-based analysis of autophagic LC3B lipidation in starved hTERT-immortalised retinal pigment epithelial (hTERT-RPE1) cells revealed impaired conversion to lipid-conjugated LC3B-II (**Fig. 1A**). In siRNA-based LC3B lipidation screens of human SNXs 1-30, we found no evidence for involvement of any other SNXs during starvation-induced autophagy (three independent tests in hTERT-RPE1 cells; data not shown). This panel included SNX18, a SNX-BAR previously shown to regulate autophagy in an imaging-based autophagy screen (Knaevelsrud et al., 2013). The lack of an effect on LC3 lipidation following treatment with SNX18 siRNA in our hands can probably be attributed to cell-type differences, assay sensitivity (imaging vs. immunoblotting), and/or siRNA efficiency. Consistent with the immunoblotting data for *SNX4* silencing (**Fig. 1A**), autophagosome puncta numbers were significantly lower in starved hTERT-RPE1 cells siRNA silenced for *SNX4* and labelled with anti-LC3B antibodies (reduced to a similar level as with *ATG5* silencing) (**Fig. 1B**).

**Figure 1:**
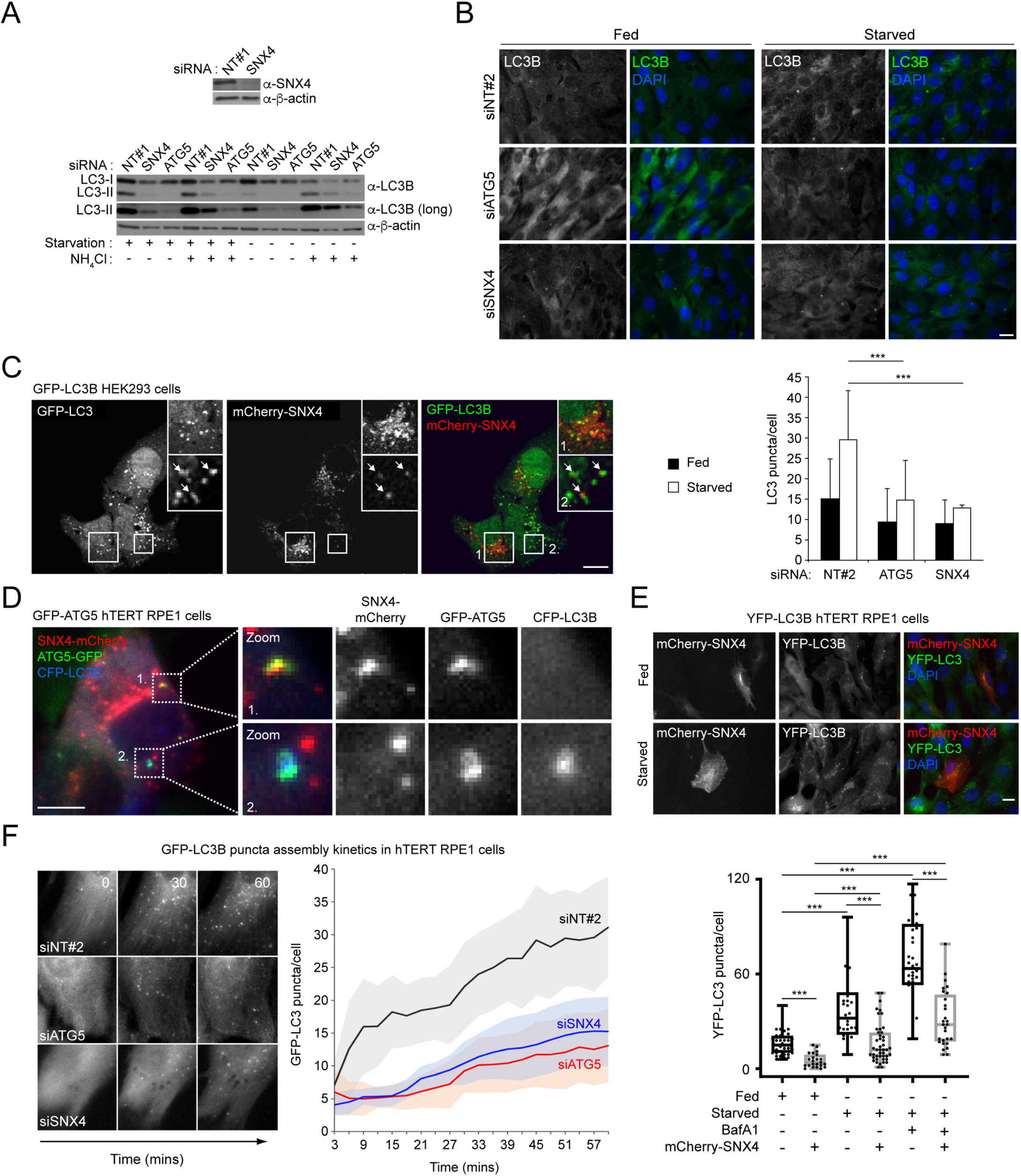
SNX4 is a positive regulator of mammalian autophagy. (A) Immunoblotting of lysates of hTERT-RPE1 cells siRNA silenced for *SNX4, ATG5* or with a non-targeting siControl (NT#1). For these experiments, hTERT-RPE1 cells were incubated for 1 h in serum and amino acid free medium (starvation) in the absence or presence of 50 mM NH_4_Cl. (B) Endogenous LC3B puncta quantitation in hTERT-RPE1 cells siRNA silenced for SNX4, ATG5 or with a non-targeting siControl (NT#2), in full nutrients (fed) and after starvation (1 h). Example images to the top; quantitation below. (C) Confocal images of GFP-LC3B HEK293 cells transiently expressing mCherry-SNX4 under starvation conditions. Coincident staining can be seen between some punctate structures. Bar = 10 µm. (D) Wide-field live-cell imaging of GFP-ATG5 hTERT RPE1 cells transiently co-expressing mCherry-SNX4 and CFP-LC3B. Examples of apparent colocalization between SNX4 and ATG5/LC3B can be seen in the zoomed areas. (E) GFP-LC3B puncta assembly kinetics during starvation in hTERT RPE1 cells treated with control siRNA (NT#2), siSNX4 or siATG5. Example fields to the left; quantitation to the right. Mean ± SE. (F) Autophagy response (LC3B puncta) in YFP-LC3B hTERT RPE1 cells transiently expressing mCherry-SNX4 in fed or starved conditions. Example fields to the top; quantitation to the bottom. ***p<0.001. Bars = 10 µm.

We have previously shown that SNX4 co-localises with peripheral endocytic membranes and the juxtanuclear RAB11-positive endocytic recycling compartment in full nutrient conditions (Traer et al., 2007). As a BAR domain protein with affinity for PtdIns(3)P, we explored the possibility that SNX4 influences autophagy via its association with autophagosomes and/or PtdIns(3)P-enriched autophagosome assembly sites. Transient expression of mCherry-SNX4 in stable GFP-LC3B HEK293 cells (Kochl et al., 2006) revealed that whilst the majority of SNX4-positive membrane structures were spatially separated from GFP-LC3B puncta, a minor fraction could be found in close proximity to GFP-LC3B labelled autophagosomes (**Fig. 1C**). Similarly, in hTERT-RPE1 cells stably expressing GFP-ATG5 (MacVicar et al., 2015), and transiently expressing mCherry-SNX4 and CFP-LC3B, we recorded incidences of SNX4-positive structures juxtaposed with GFP-ATG5-positive autophagosome assembly sites, and further examples of mCherry-SNX4 structures in close proximity to GFP-ATG5 and CFP-LC3B-positive structures likely to be forming phagophores (**Fig. 1D**), suggesting that any interaction is likely to occur during autophagosome assembly, at around the ATG8 lipidation stage.

To begin to understand how SNX4 influences autophagy, we imaged autophagosome assembly kinetics during amino acid/growth factor starvation in hTERT-RPE1 cells stably expressing GFP-LC3B (**Fig. 1E**). We compared control, *ATG5*, and *SNX4* siRNA silenced cells, assessing cumulative GFP-LC3B puncta numbers without inclusion of lysosomal blocking reagents. The kinetics of GFP-LC3B puncta assembly were clearly altered when *SNX4* was silenced, with puncta formation rates decreased to a level that was comparable with *ATG5*-silenced cells (**Fig. 1E**). This suggested that the reduction in stimulated LC3B-positive autophagosome numbers observed at steady state in fixed cells (**Fig. 1B**) was unlikely to be due to increased LC3B turnover due to enhanced autophagic flux. As overexpression of SNX18 enhanced LC3 lipidation without influencing autophagic flux (Knaevelsrud et al., 2013), we tested the effects of mCherry-SNX4 overexpression on the autophagy response in hTERT-RPE1 cells stably expressing YFP-LC3B, counting basal and starvation-induced LC3B puncta numbers in mCherry-SNX4-positive cells (**Fig. 1F**). Basal (fed state) LC3B numbers in were significantly lower in mCherry-SNX4 overexpressing cells when compared with neighbouring, untransfected cells, and this pattern was repeated following starvation in the absence of presence of the vacuolar H^+^-ATPase inhibitor, Bafilomycin A1 (BafA1) (**Fig. 1F**). This suggests that SNX4 overexpression impairs autophagosome assembly, rather than increasing the rate of lysosomal flux. Steady state numbers of GFP-ATG5-positive assembly sites were not affected by SNX4 overexpression (**Fig. S1**). These data suggest that the balance of expression of different autophagy-influencing SNX-BARs effects autophagy in different ways; whilst SNX18 overexpression enhanced LC3 lipidation (Knaevelsrud et al., 2013), elevated SNX4 expression clearly suppressed the starvation-mediated autophagy response at the level of LC3B puncta formation.

The observed differences in autophagy responses in cells overexpressing SNX4 might relate to changes in the structure/function of the endocytic compartment. To examine this, we generated hTERT-RPE1 cell-lines stably overexpressing GFP-SNX4. In these cells, the number of distinct EEA1-positive early endocytic structures was significantly lower than seen in control, GFP-expressing hTERT-RPE1 cells under both fed and starvation conditions (**Fig. S2A**). Analysis of CD63-positive lysosomes revealed significantly more lysosomes in fed GFP-SNX4 expressing hTERT-RPE1 cells, but a clear absence of induced CD63 puncta increases following amino acid/growth factor starvation when compared to control GFP-expressing cells (**Fig. S2B**). This suggests that SNX4 overexpression upsets the balance of membrane trafficking within the endolysosomal compartment. To rule out the possibility that SNX4 overexpression disturbs endocytic organelle properties and autophagosome assembly kinetics by competing for and/or blocking available PtdIns(3)P on endosomes and autophagosome assembly sites, we analysed LC3B puncta numbers in cells overexpressing a similar PtdIns(3)P-binding motif, mCherry-2xFYVE (**Fig. S3**). No differences in LC3B puncta numbers were recorded in mCherry-2xFYVE expressing cells in fed or starved conditions or when treated with the mTORC1/2 inhibitor, AZD8055, when compared to untransfected controls (**Fig. S3**). Together, these data suggest that reduced autophagosome assembly/LC3B lipidation kinetics caused by changes in the levels of cytosolic SNX4 in nutrient starved cells (by overexpression or by siRNA silencing) may be linked to altered endomembrane properties. This prompted us to further explore further the endosomal biology of SNX4 and its relationship with the autophagy regulatory system.

### SNX4 displays a restricted pattern of interactions with two other SNX-BAR family members, SNX7 and SNX30

Within the SNX-BAR family, there is strong evidence for a restricted pattern of BAR domain-mediated dimerisations, leading to the formation of specific SNX-BAR homo- and heterodimers (Cullen, 2008; Simonetti et al., 2017; van Weering et al., 2012; van Weering et al., 2010). Previous analysis using overexpressed proteins indicated that SNX4 forms weak homodimers and relatively stable heterodimers with the SNX-BARs, SNX7 and SNX30 (van Weering et al., 2012). To begin to assess whether SNX4 influences autophagy in homodimeric form or in heterodimeric complex with another SNX-BAR, we carried out directed yeast two-hybrid screens, probing the interactions of full-length SNX4 with alternate SNX-BARs (SNX7, SNX8 and SNX30, and the ESCPE-1 SNX-BARs, SNX1, SNX2, SNX5, SNX6, and SNX32) (**Fig. 2A**). SNX4 did not form any detectable associations with ESCPE-1 SNX-BARs or with SNX8; however, a weak interaction with itself and strong interactions with both SNX7 and SNX30 were observed (**Fig. 2A**), consistent with previous in vitro pull-down analysis (van Weering et al., 2012). A limitation of interactions studies requiring the overexpression of one or other putative partner protein concerns the forcing of interactions that may not be physiologically relevant in vivo. We therefore sort further information on the specificity of interactions between SNX4, SNX7 and SNX30 through immunoprecipitation of endogenous proteins (**Fig. 2B**). Confirming previously published data describing the restricted pattern of heterodimeric interactions between SNX1:SNX5 and SNX1:SNX6 (Wassmer et al., 2007; Wassmer et al., 2009), immunoprecipitates of SNX1 were positive for endogenous SNX5 and SNX6 (**Fig. 2B**). Within these immunoprecipitates, we failed to detect endogenous SNX30, SNX9, SNX8, SNX7 or SNX4 (**Fig. 2B**). Conversely, immunoprecipitates of endogenous SNX4 were characterised by the presence of endogenous SNX7 and SNX30, and the lack of any detectable retromer SNX-BARs, or SNX9 and SNX8 (**Fig. 2B**). More specifically, within immunoprecipitates of endogenous SNX7 and SNX30, the only other sorting nexin detected in each case was SNX4 (**Fig. 2B**). Overall, these data clearly establish that, at the level of endogenous protein expression, SNX4 forms the core of two distinct heterodimeric complexes; SNX4:SNX7 and SNX4:SNX30.

**Figure 2:**
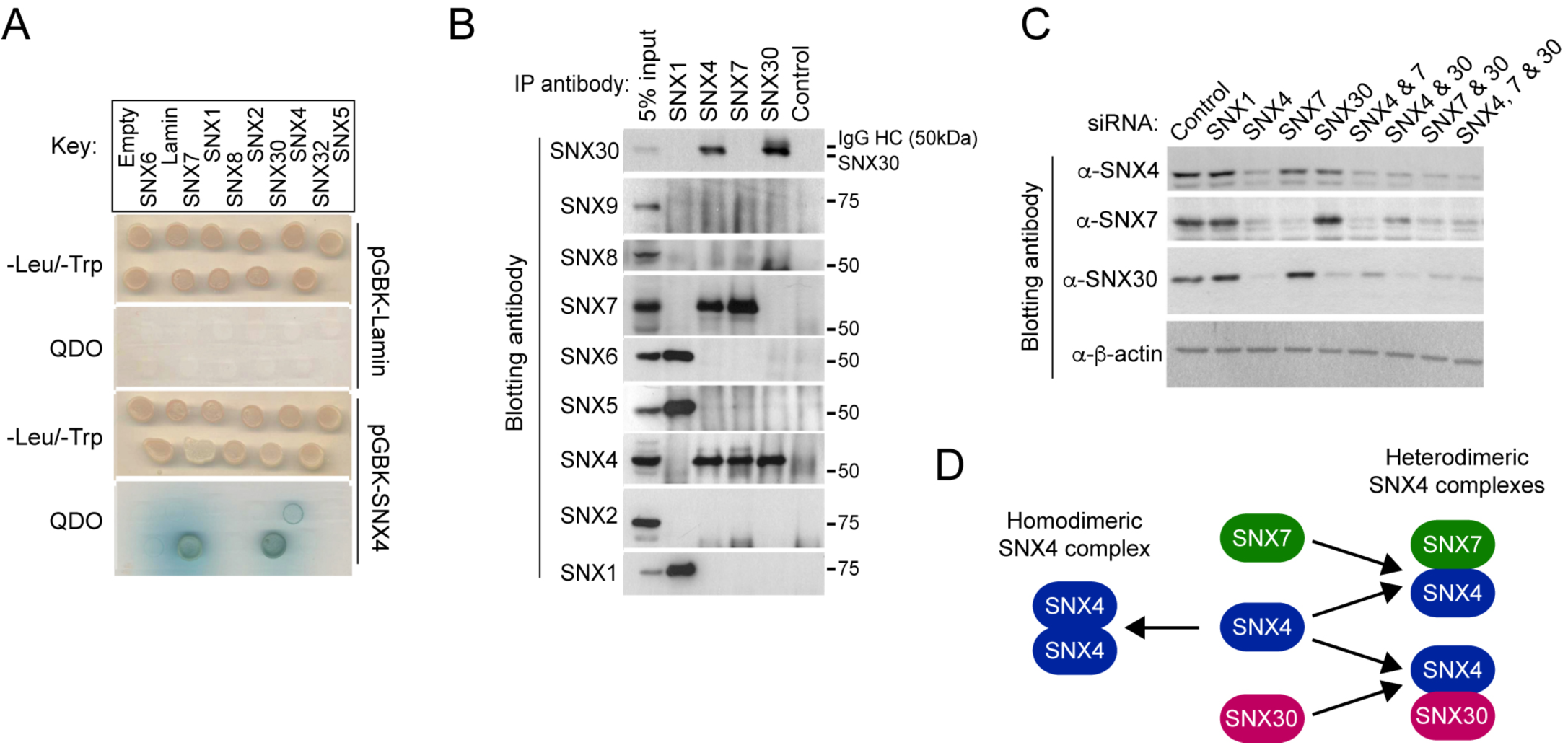
SNX4 forms the core component of 2 distinct SNX-BAR heterodimers. (A) Directed yeast 2-hybrid test for SNX-BAR interactions. Lamin is included as a negative control. (B) Native immunoprecipitations of SNX1, SNX4, SNX7 and SNX30 in HeLa cells. Immunoprecipitates were blotted for the SNXs shown. (C) siRNA silencing of SNX1 and SNX4, SNX7, SNX30 alone and in combination in HeLa cells. Lysates were immunoblotted using the antibodies shown. (D) Schematic of the likely dimerisations that exist between SNX4/SNX7/SNX30 SNX-BAR pairings.

For the mammalian ESCPE-1 SNX-BARs, it is well documented that siRNA suppression of the expression of SNX1 and SNX2 destablises SNX5 and SNX6, leading to their degradation (Carlton et al., 2004; Carlton et al., 2005; Simonetti et al., 2017; Wassmer et al., 2007). As this could have consequences for our interpretation of autophagy in *SNX4* silenced cells, we considered whether a similar scenario exists for the SNX4:SNX7 and SNX4:SNX30 dimers: if SNX4 is the core component, does its suppression also lead to a loss of SNX7 and/or SNX30? Indeed, siRNA suppression of SNX4 protein expression caused a clear decrease in the levels of both SNX7 and SNX30 (**Fig. 2C**). In contrast, suppression of the retromer component SNX1 had no discernible effect on SNX4, SNX7 or SNX30 (**Fig. 2C**). Interestingly, effective silencing of SNX7 expression resulted in a small but clearly detectable drop in SNX4 protein levels, while the expression of SNX30 appeared unaffected (**Fig. 2C**). Likewise, strong suppression of SNX30 expression again induced a detectable decrease in SNX4, but this had no effect on SNX7 expression (**Fig. 2C**). Taken together, these data are consistent with SNX4 forming the core of two distinct SNX4:SNX7 and SNX4:SNX30 complexes (**Fig. 2D**). Thus, upon SNX4 suppression, the loss of the core component destabilises the other constituents, while upon suppression of an individual complex specific component, such as SNX7, the presence of the core SNX4 component allows the stabilisation of the SNX4:SNX30 complex (and vice versa under conditions of SNX30 suppression). Indeed, as one would predict from such a model, dual suppression of SNX7 together with SNX30 led to a pronounced loss in the levels of SNX4 (**Fig. 2C**).

### SNX4 co-localises with SNX7 and SNX30 predominantly on early endosomes

Previous studies have established that SNX4 is targeted to early endosomes via association of its PX domain with PtdIns(3)P (Leprince et al., 2003; Skanland et al., 2009; Skanland et al., 2007; Teasdale et al., 2001; Traer et al., 2007). While the phosphoinositide-binding characteristics of SNX30 have yet to be described, SNX7 has been shown to associate specifically with PtdIns(3)P (Xu et al., 2001). To examine whether SNX7 and SNX30 also associate with PtdIns(3)P-enriched early endosomes, we used lentiviruses to express GFP- or mCherry-tagged full length human SNX7 or SNX30 at a level that did not cause noticeable remodelling of endosomal membranes (Carlton et al., 2004; Cozier et al., 2002). Unlike the situation when expressing GFP-tagged SNX4 (Traer et al., 2007), lentiviral transduction of HeLa cells with lentiviruses encoding GFP-SNX7 or GFP-SNX30 alone resulted in relatively weak levels of expression, and in those cells where a signal could be observed, both gave predominantly cytosolic staining patterns with some evidence of punctate staining (**Fig. S4**). Interestingly, when these same viruses were used to co-transduce HeLa cells alongside a virus encoding mCherry-SNX4, clear co-localisation of mCherry-SNX4 and GFP-SNX7/GFP-SNX30 was readily observed on puncta that were dispersed throughout the cytoplasm (**Fig. 4A, B; Fig. S4A, B**). Based on the biochemical evidence that SNX7 and SNX30 are unable to establish stable homodimers (**Fig. 2**), and given that dimerization is a prerequisite for assembly of a functional membrane binding BAR domain (Peter et al., 2004), we interpret these data to mean that in the absence of co-expression with SNX4, SNX7 and SNX30 exist as unstable monomers that have insufficient affinity for PtdIns(3)P to attain steady-state endosomal association. Indeed, the importance of the combined, co-incidence membrane binding activities of the PX and BAR domains is well established in this context (Carlton et al., 2004; Traer et al., 2007). Although the colocalisation of SNX7 and SNX30 with SNX4 was consistent with an association with early endosomes, we addressed this directly by co-transducing HeLa cells with lentiviruses encoding GFP-SNX7 or GFP-SNX30 together with a lentivirus encoding for FLAG-tagged SNX4, and counter stained with a variety of early and late endosomal markers (**Fig. 3C, D**). Confocal imaging revealed that in the presence of FLAG-SNX4, both GFP-SNX7 and GFP-SNX30 decorated early endocytic structures that partially overlapped with EEA1 and SNX1, but did not correlate with APPL1 or CD63 (very early endosomal and late endosomal/lysosomal markers, respectively), suggesting that these SNX-BARs associate with intermediate stage endocytic structures (**Fig. 3C, D**), consistent with the previously reported distribution of SNX4 (Traer et al., 2007).

**Figure 3:**
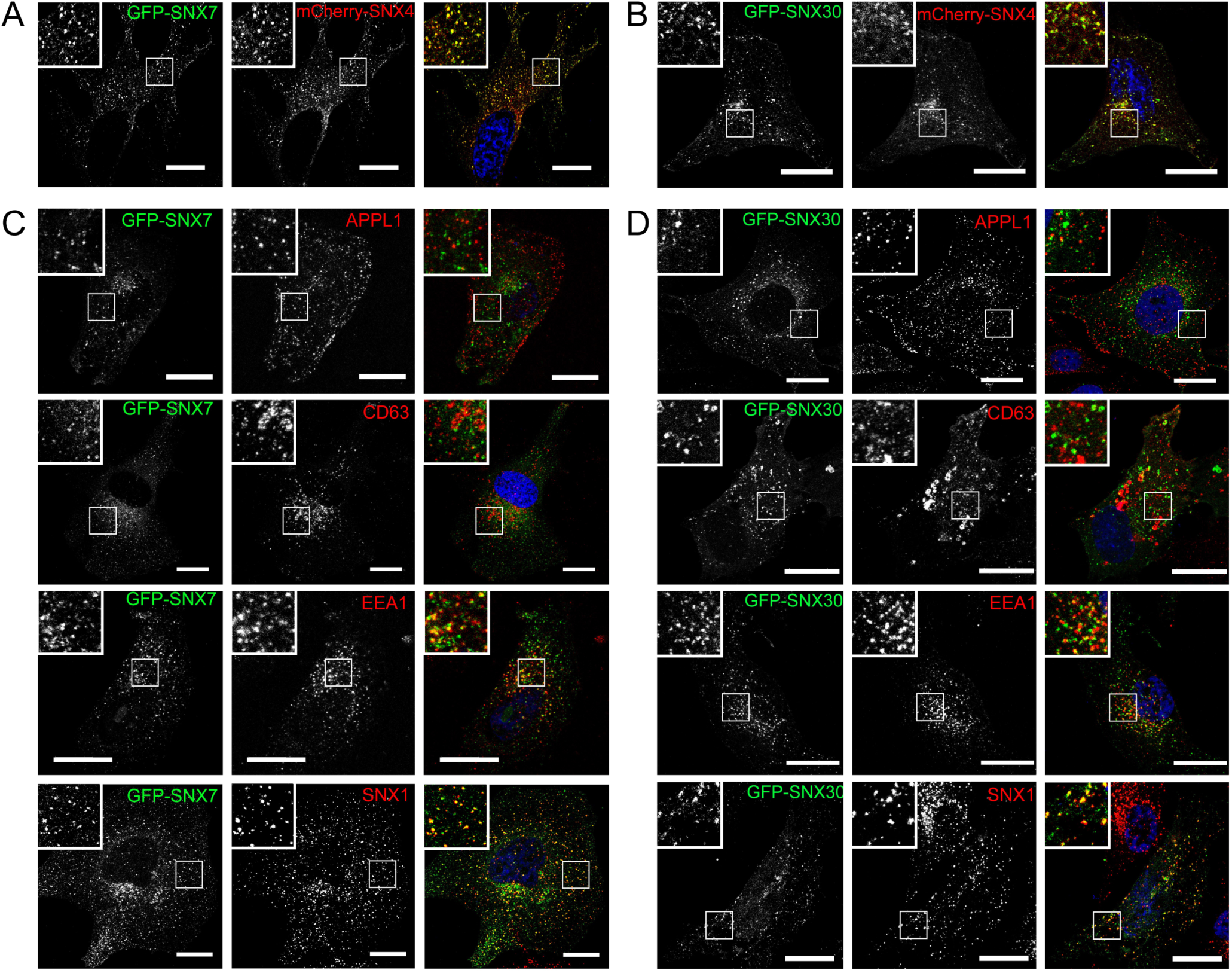
Subcellular localisation of SNX4, SNX7 and SNX30. (A, B) Confocal images of HeLa cells expressing GFP-SNX7 (A) or GFP-SNX30 (B) with mCherry-SNX4. The requirement for SNX4 co-expression is further assessed in **Fig. S3**. (C, D) Analysis of GFP-SNX7 (C) and GFP-SNX30 (D) endosomal targeting in HeLa cells co-expressing FLAG-SNX4 (not shown) and counterstained for various endocytic markers (APPL; CD63; EEA1; SNX1). Bar = 10 µm.

**Figure 4:**
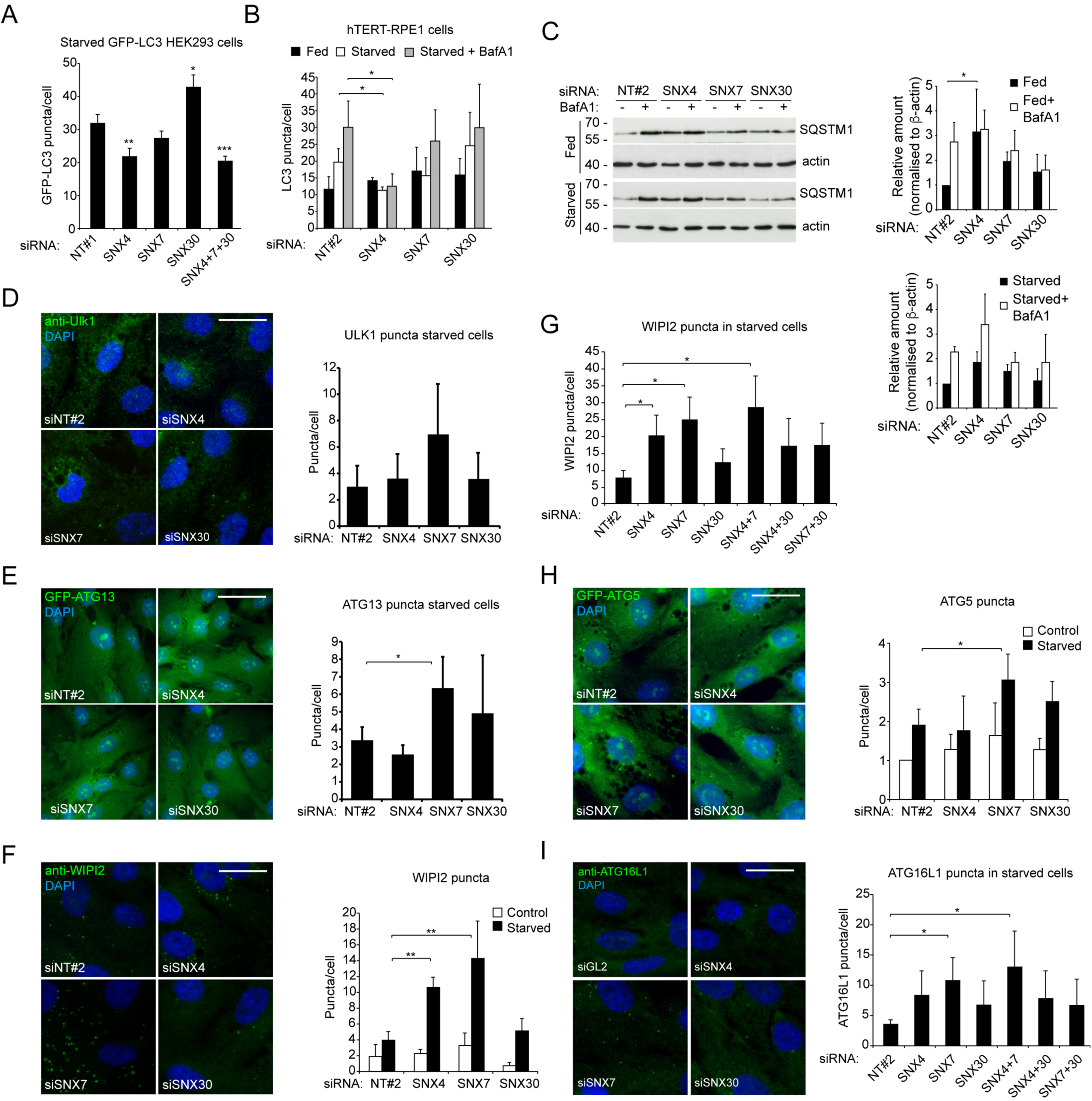
The SNX4:SNX7 heterodimer is required for efficient autophagy. (A) Autophagy responses (LC3B puncta) measured in starved GFP-LC3B HEK293 cells treated with SNX4/SNX7/SNX30 siRNA alone or in combination. Mean ± SE. (B) Autophagic flux analysis in hTERT-RPE1 cells silenced for SNX4, SNX7 or SNX30, and labelled for endogenous LC3B. Mean ± SD. (C) p62/SQSTM1 turnover assay in hTERT-RPE1 cells silenced for SNX4, SNX7 or SNX30. Example blots to the left; quantitation to the right. Mean ± SD. (D-I) Puncta analysis in SNX4, SNX7 and SNX30 siRNA treated hTERT-RPE1 cells labelled with antibodies or stably expressing GFP fusions of autophagy markers, with example images and quantitation shown as follows: (D) Endogenous ULK1 puncta in starved cells; (E) GFP-ATG13 puncta in starved GFP-ATG13 stable cells; (F) Endogenous WIPI2 puncta in fed (control) and staved cells; (G) Endogenous WIPI2 puncta in starved cells treated with combinatorial siRNAs; (H) GFP-ATG5 puncta in fed (control) or starved GFP-ATG5 stable cells; (I) Endogenous ATG16L1 in cells treated with combinatorial siRNAs. Non-targeting siRNA NT#1 is included as control in (A); NT#2 in (B-I). Bars = 20 µm. Parts D-I show means ± SD; *p<0.05; **p<0.01; ***p<0.001.

### Autophagic flux defects in SNX4/7/30 suppressed cells

To test for the existence of an autophagy-specific SNX4-containing SNX-BAR heterodimer, we measured whether siRNA suppression of either *SNX7* or *SNX30* caused defects at the level of LC3B puncta formation. As expected, in GFP-LC3B HEK293 cells, *SNX4* suppression again caused a significant reduction in LC3B puncta numbers during starvation, whereas suppression of SNX7 levels did not alter the autophagy response (**Fig. 4A**). Interestingly, in these cells, we recorded a significant increase in GFP-LC3B puncta numbers after *SNX30* silencing (**Fig. 4A**). Possible explanations include: (i) the SNX4:SNX30 heterodimer acting to restrict autophagosome assembly; (ii) SNX30 suppression indirectly enhancing LC3B lipidation by biasing assembly of an autophagy-enhancing SNX4:SNX7 heterodimeric complex; or (iii) reduced autophagic flux when SNX30 is suppressed. The latter appeared unlikely as combinatorial siRNA suppression of all 3 SNXs effectively phenocopied siRNA silencing of *SNX4*, with a significant decrease in LC3 puncta recorded (**Fig. 4A**).

To assess the impact of reduced SNX4/7/30 levels on autophagy in cells expressing only endogenous LC3B—and to clarify its impact on autophagic flux (**Fig. 4A**)—we treated hTERT-RPE1 cells with the appropriate siRNA oligonucleotides before starving them in the absence or presence of BafA1 (**Fig. 4B**). A significant reduction in LC3B puncta numbers was recorded in *SNX4*-silenced hTERT-RPE1s starved in the absence and presence of BafA1, suggesting that the reduced autophagy response was most likely due to the failure to assemble LC3B-positive autophagosomes, rather than an increase in the rate of flux (i.e. LC3 puncta turnover) (**Fig. 4B**). Basal autophagy levels were similar between conditions (**Fig. 4B**). In these assays, the starvation-induced autophagy response in *SNX7*-silenced hTERT-RPE1 cells was consistently lower than in control and in *SNX30*-silenced cells, although this was not statistically significant (**Fig. 4B**). Finally, in contrast to GFP-LC3B expressing HEK293 cells, silencing of *SNX30* had no impact on the numbers of endogenous LC3B puncta in hTERT-RPE1s (**Fig. 4B**). This probably reflects cell-type specific differences in the relative balance of SNX4-containing heterodimers and/or the efficiency of siRNA silencing between cell-lines. To assess the consequences of SNX4/7/30 suppression on autophagy cargo flux, we carried out p62/SQSTM1 turnover assays in hTERT-RPE1 cells starved in the absence or presence of BafA1. In *SNX4*-silenced cells under basal conditions, p62/SQSTM1 levels were significantly higher than in controls, and incubation in the presence of BafA1 did not increase p62/SQSTM1 levels, indicative of a block in autophagic flux in full nutrients (**Fig. 4C**). Interestingly, in hTERT-RPE1 cells suppressed for either SNX7 or SNX30, the anticipated increase in p62/SQSTM1 levels following BafA1 treatment was also absent, although under these conditions, basal p62/SQSTM1 levels were not significantly different from controls (**Fig. 4C**). It was also notable that in *SNX4*-silenced hTERT-RPE1 cells, p62/SQSTM1 levels did increase following addition of BafA1 during starvation, although this was not statistically significant (**Fig. 4C**). This suggests that when SNX4 levels are reduced, autophagic p62/SQSTM1 turnover can still occur, but with reduced efficiency. Taken together, the interactions and functional data show that SNX4 can exist either as a weak homodimer, or as the core component of 2 distinct heterodimeric SNX-BAR complexes; SNX4:SNX7 and SNX4:SNX30. Which if any of these complexes acted selectively to regulate the autophagy process remained unclear.

### SNX4:SNX7 is an autophagy SNX-BAR heterodimer required during early stages of autophagosome assembly

Dynamic imaging studies have provided a model of quasi-hierarchical recruitment of autophagy regulators to the autophagosome assembly site, with both forward and reverse reinforcement interactions between key players (e.g. (Karanasios and Ktistakis, 2015)). To determine how the SNX4 homodimer and/or SNX4:SNX7/SNX4:SNX30 heterodimers influence autophagosome formation, the recruitment and retention of key autophagy regulators at the autophagosome assembly site were assessed in hTERT-RPE1 cells by fluorescence microscopy (**Fig. 4D-I**). In this analysis, both reduced and increased autophagosome assembly site numbers can indicate a kinetic block in autophagosome assembly, when assessed alongside other tests for autophagy (e.g. LC3 puncta numbers; p62/SQSTM1 turnover). For example, reduced numbers of assembly sites can be consistent with either a block in early signalling and/or the failure to recruit/stabilise early mediators at the autophagosome assembly site; by contrast, elevated numbers of assembly site foci can indicate enhanced autophagy signalling, or assembly site stalling (as also seen in cells depleted for SNX18 (Knaevelsrud et al., 2013)).

We began by analysing markers of the ULK1 kinase complex using anti-ULK1 antibodies and a stable GFP-ATG13 hTERT-RPE1 cell-line following amino acid/growth factor withdrawal (**Fig. 4D, E**). Whilst there were no differences in the steady-state numbers of ULK1 and GFP-ATG13 puncta in SNX4- and SNX30-suppressed cells compared to controls, starvation-induced ULK1 and GFP-ATG13 puncta numbers were elevated in SNX7-suppressed cells (although this was only statistically significant for GFP-ATG13 [**Fig. 4D, E**]). These data established for the first time a possible role for SNX7 in the regulation of autophagosome assembly dynamics, supporting the earlier hint that its suppression might be restricting LC3B lipidation in the same cell-type (**Fig. 4B**). We next analysed subsequent stages of assembly site maturation—namely, PtdIns(3)P enrichment, and the recruitment/retention of the ATG8 lipidation machinery—using antibodies against WIPI2 and ATG16L1, and using the GFP-ATG5 stable hTERT-RPE1 cell-line (MacVicar et al., 2015) (**Fig. 4F-I**). In common with the GFP-ATG13 data (**Fig. 4E**), we recorded significantly higher starvation-induced WIPI2 puncta numbers in SNX7-silenced cells, confirming that autophagosome assembly is indeed sensitive to SNX7 levels in human cells (**Fig. 4F, G**). Starvation-induced WIPI2 puncta numbers were also significantly higher in cells suppressed for SNX4 (**Fig. 4F, G**); however, GFP-ATG5 puncta numbers did not differ significantly from controls in SNX4-suppressed cells (**Fig. 4H**).

To tease out the possible contributions of the different SNX4-containing heterodimers, we next analysed the effects of combinatorial suppression of SNX4/7/30 in starved hTERT-RPE1 cells labelled with WIPI2, and with another marker for the ATG8 lipidation machinery, ATG16L1 (**Fig. 4G, I**). Analysis of ATG16L1-labeled cells revealed the now expected pattern of elevated ATG16L1 puncta numbers in SNX7-suppressed cells, and a clear, but not statistically significant increase in ATG16L1 puncta in SNX4-suppressed cells (**Fig. 4I**). The lack of an additive effect on ATG16L1 (**Fig. 4G**) and WIPI2 (**Fig. 4I**) puncta numbers following SNX4/7 co-suppression indicated that these SNX-BARs might act in concert to regulate autophagy. Of note, we found that both ATG16L1 and WIPI2 puncta numbers did not diverge significantly from control levels when cells were co-suppressed for SNX4 with SNX30, and for SNX7 with SNX30 (**Fig. 4G, I**). Evidently, a shift in the relative levels of the cognate SNX4 dimers, due to siRNA depletion of individual components (see also **Fig. 3C**), can alter the autophagy response in complex ways.

Puncta analysis suggested that there are likely to be kinetic differences in the recruitment and/or retention of key autophagy markers at the isolation membrane during early autophagosome formation in SNX4/7 suppressed cells. To further assess where defects arose, we carried out co-localisation analysis of fixed GFP-ATG5 hTERT-RPE1 cells labelled with anti-WIPI2 and anti-ATG16L1 antibodies (**Fig. 5A**). Each of these markers is recruited to the isolation membrane at a similar stage in advance of ATG8 lipidation. Nevertheless, we found that although there were no changes in ATG16L1/ATG5 co-localisation when cells were suppressed for SNX4, SNX7 or SNX30 (**Fig. 5B**), co-localisation between WIPI2 and ATG5 was lower in SNX4- and SNX7-suppressed cells (although this was significant only for SNX7; **Fig. 5C**). This suggests that effective ATG5 recruitment and/or retention at WIPI2-positive PtdIns(3)P early autophagic structures depends upon the presence of the SNX4:SNX7 heterodimer. The absence of any specific defect at the level of ATG5 puncta numbers following SNX4 suppression (**Fig. 4H**) was surprising; however, the impact of *SNX4* silencing on the kinetics of assembly site assembly/disassembly could be revealed during live imaging of the GFP-ATG5 hTERT-RPE1 cell-line (**Fig. 5D-F**). In control siRNA depleted cells, and in cells depleted for either SNX7 or SNX30, average GFP-ATG5 puncta lifetime was on average ∼2.5 mins (**Fig. 5D, E**). By contrast, in *SNX4*-silenced cells, average GFP-ATG5 length was significantly shorter at ∼1.75 mins (**Fig. 5D**), with the distribution of GFP-ATG5 puncta lifetimes clearly altered (**Fig. 5E**). Furthermore, time-resolved comparisons revealed that in *SNX4*-silenced cells, GFP-ATG5 fluorescence intensities were consistently lower than in control cells—although this was not statistically significant at any individual time-point (**Fig. 5F**). Together these data suggest that ATG5 recruitment and/or turnover kinetics are altered in SNX4-suppressed cells (**Fig. 5C-F**), and that cells most likely compensate for this by upregulating autophagosome assembly sites, meaning that steady state GFP-ATG5 puncta numbers appear similar to control cells (**Fig. 4H**).

**Figure 5:**
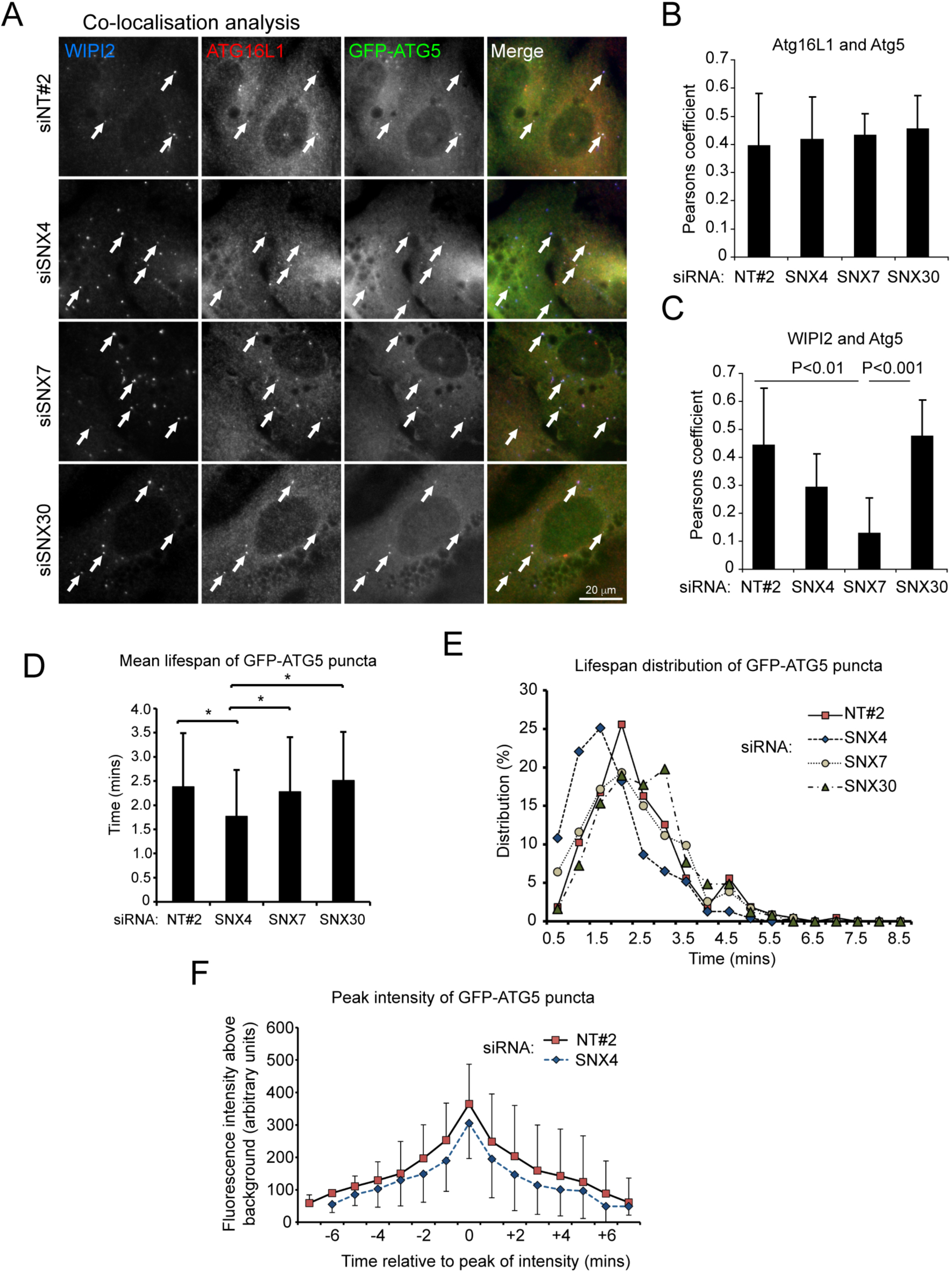
SNX4/SNX7 depletion impacts on autophagy at the ATG5 recruitment/retention stage. (A-C) Co-localisation analysis of ATG16L1 with ATG5, and WIPI2 with ATG5 in GFP-ATG5 stable hTERT-RPE1 cells. (A) Example images; arrows show examples of colocalising puncta. (B-C) Pearson’s coefficient colocalisation between (B) ATG16L1 and ATG5, and (C) WIPI2 and ATG5. Mean ± SD; bar = 20 µm. (D) GFP-ATG5 puncta lifetime in stable hTERT-RPE1 cells silenced for SNX4, SNX7 or SNX30. Mean ± SD. (E) GFP-ATG5 puncta lifespan distribution in stable hTERT-RPE1 cells silenced for SNX4, SNX7 or SNX30. (F) Peak GFP-ATG5 puncta fluorescence intensity analysis in control (NT#2) and SNX4 siRNA-treated GFP-ATG5 stable hTERT-RPE1 cells. *p<0.05; **p<0.01; ***p<0.001.

### ATG9A trafficking is defective in SNX4 CRISPR knockout (KO) HeLa cells

Our siRNA-based results highlighted how the SNX4:SNX7 heterodimer acts during autophagosome formation at the ATG5 recruitment stage. During autophagosome assembly, the ATG12∼ATG5 conjugate is recruited via binding to ATG16L1, with additional membrane binding capability conferred by ATG5 itself (Romanov et al., 2012). To determine how the SNX4:SNX7 heterodimer influences this step, we used CRISPR-Cas9 to generate HeLa cell-lines edited to eliminate SNX4 expression. Several clones were produced that showed reduced or absent SNX4 levels, and these were assessed for relative SNX7 and SNX30 expression (**Fig. S5**). We selected clone “A” for detailed analysis as these cells showed depleted SNX7 and SNX30 alongside an absence of SNX4 (**Fig. S5**; **Fig. 6A**). Analysis of the autophagy response in these cells following application of AZD8055 (2 hours) revealed a significant suppression of LC3B lipidation, and corresponding reduced numbers of LC3B-positive autophagosomes in SNX4 KO cells in the absence and presence of BafA1 (**Fig. 6B, C**). Interestingly, in contrast to *SNX4* siRNA suppressed hTERT-RPE1 cells in which WIPI2 puncta levels were significantly higher than in wild-type cells during autophagy stimulation (**Fig. 4F, G**), WIPI2 puncta numbers in SNX4 KO cells did not differ from wild-type, with or without autophagy stimulation (**Fig. 6C**). This variability might be due to the different cell-types tested (hTERT-RPE1 cells vs. HeLa cells) and/or because of compensatory pathways emerging in the SNX4 KO cells. Interestingly, WIPI2 puncta were found to be lower in SNX18 KO cells than in wild type cells following autophagy stimulation (Soreng et al., 2018), suggesting that these SNX-BARs influence different stages of autophagosome assembly. To assess autophagic flux, we generated mCherry-GFP-LC3B stable wild-type and SNX4 KO cells, and assessed autophagosome (red/green) and autolysosome (red only as GFP is quenched at acidic pH) puncta numbers following 2 hours AZD8055 treatment in the absence or presence of BafA1 (**Fig. 6D, E**). Green/red-positive (yellow) autophagosome numbers were significantly fewer in the SNX4 KO cells than in controls under all treatments conditions except the basal state (i.e. treated with BafA1, AZD, or AZD + BafA1) (**Fig. 6D, E**). This suggests that autophagic flux is intact in SNX4 KO cells, but the efficiency of both assembly and flux is impaired. In fed SNX4 KO cells, LC3B-positive autolysosomes (red) were significantly more abundant than in control cells (**Fig. 6E**). This unexpected finding suggests that basal autophagic flux may be less efficient in the SNX4 KO cells, although the defect was not retained during autophagy stimulation (**Fig. 6E**; the effective reduction in LC3-positive puncta numbers in SNX4 KO cells after autophagy stimulation is possibly being caused by increased lysosomal clustering and/or fusion). As further evidence that loss of SNX4 affected autophagosome assembly, rather than lysosomal fusion (flux), we analysed colocalization between red and green puncta under the same conditions (**Fig. 6F**). As expected, colocalization increased when lysosomes were inhibited using BafA1 in control and AZD-treated cells (as a result of impaired GFP quenching), but no differences were observed in any single condition between control and SNX4 KO cells (**Fig. 6F**). Importantly, in SNX4 KO cells, transient expression of SNX4-mCherry was sufficient to rescue the autophagy defect at the level of LC3B puncta formation (**Fig. 6G**).

**Figure 6:**
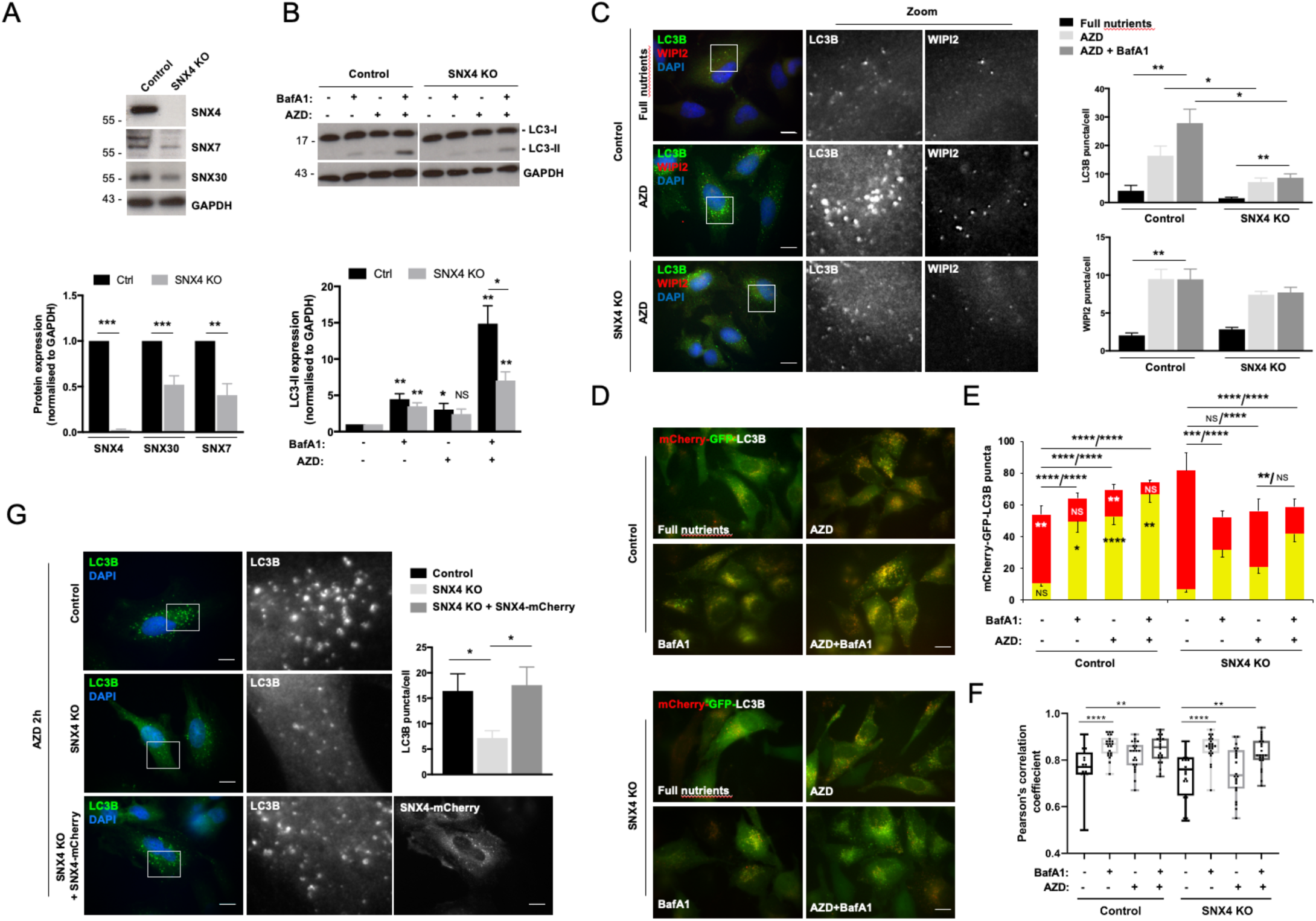
A SNX4 CRISPR KO HeLa cell-line is autophagy deficient. (A) Immunoblotting analysis of SNX4, SNX7 and SNX30 levels in parental cells and the SNX4 CRISPR KO line. Example blots above; quantitation below. (B) Immunoblotting-based LC3B lipidation analysis in the SNX4 CRISPR KO line treated with AZD8055 in the absence or presence of BafA1. Example blots above; quantitation below. (C) LC3B and WIPI2 puncta analysis in the SNX4 CRISPR KO line treated with AZD8055 in the absence or presence of BafA1. Example images to the left; quantitation to the right. Bar = 10 µm. (D) Example live images of control and SNX4 KO HeLa cells stably expressing mCherry-GFP-LC3B, treated with BafA1, AZD8055, or BafA1/AZD8055. Bar = 10 µm. (E) mCherry-GFP-LC3B flux assessment. Means ± of ∼20 cells per condition, imaged live. Asterisks superimposed upon the control cell bars represent statistical comparisons between control and SNX4 KO cells (one-way ANOVA); asterisks above the bars represent indicated pairwise comparisons for green/red analysis (Welch’s t test; NS = not significant). (F) Colocalisation analysis (Pearson’s) of mCherry- and GFP-positive LC3B puncta in live control and SNX4 KO HeLa cells stably expressing mCherry-GFP-LC3B (Kruskal-Wallis non-parametric ANOVA). (G) Rescue of the autophagy (LC3B puncta) defect in SNX4 CRISPR KO HeLa cells transiently expressing mCherry-SNX4. Bar = 10 µm. *p<0.05; **p<0.01; ***p<0.001; ****p<0.0001.

The multi-pass transmembrane protein ATG9A traffics through the endocytic network to establish a perinuclear RAB11/TGN-associated compartment from where it can be rapidly mobilised upon autophagy stimulation to form peripheral membrane pools required for efficient autophagosome biogenesis (for a recent discussion, see (Shatz and Elazar, 2019)). Since SNX18 has been found to coordinate ATG9A redistribution during autophagy via the actions of dynamin-2 (Soreng et al., 2018), we tested whether SNX4 also influences ATG9A trafficking during autophagy stimulation. We carried out immunofluorescence analysis of ATG9A localisation in cells counterstained for the Golgi marker, GM130 (**Fig. 7A, B**), or transferrin receptor (TfR) (**Fig. 7C**). In wild-type HeLa cells, ATG9A steady state localisation comprised a prominent membrane pool focussed in the Golgi region, with additional dispersed vesicular profiles (**Fig. 7A**). As reported by others (e.g. (Soreng et al., 2018; Young et al., 2006)), autophagy stimulation (AZD8055, 2 hours) caused further dispersal of the ATG9A Golgi pool (**Fig. 7A, B**), while TfR localisation remained largely unchanged (**Fig. 7C**). At steady state (full nutrients; without AZD8055), ATG9A showed a significantly stronger colocalisation with GM130 in SNX4 KO cells when compared to controls (**Fig. 7A**), and whilst in both cell-types ATG9A became more vesicular/dispersed following 2 hours AZD8055 treatment, ATG9A remained more closely associated with the Golgi region in the SNX4 KO cells, suggesting defective mobilisation of the ATG9A Golgi pool following autophagy stimulation (**Fig. 7A**). Importantly, the ATG9A redistribution defect could be rescued by transient expression of SNX4-mCherry in SNX4 KO cells (**Fig. 7B**). Interestingly, we noted some important differences in ATG9A behaviour between SNX4 KO and SNX18 KO cells (Soreng et al., 2018) with respect to TfR trafficking. Although ATG9A colocalisation with TfR was enhanced by SNX18 depletion, with TfR becoming more juxtanuclear in the absence of SNX18 (Soreng et al., 2018), in SNX4 KO cells, ATG9A colocalisation with TfR was significantly lower than in control cells, and changed little following AZD8055 treatment (**Fig. 7C**). Finally, and despite the lack of evidence for an interaction between SNX4 and ATG9A at the biochemical level (this was also true for other autophagy proteins; **Fig. S6**), SNX4-mCherry transiently expressed in the SNX4 KO background showed a strong colocalisation with ATG9A (**Fig. 7D**), suggesting that a sub-fraction of each resides in the same endomembrane compartment. Together, these data suggest that SNX4 contributes to the steady state localisation of ATG9A, and that SNX4 is required for efficient ATG9A peripheral redistribution upon autophagy stimulation to enable efficient autophagy responses. Our data are consistent with the SNX4:SNX7 autophagy SNX-BAR heterodimer contributing to the control of ATG9A trafficking to the autophagosome assembly site during autophagy stimulation.

**Figure 7:**
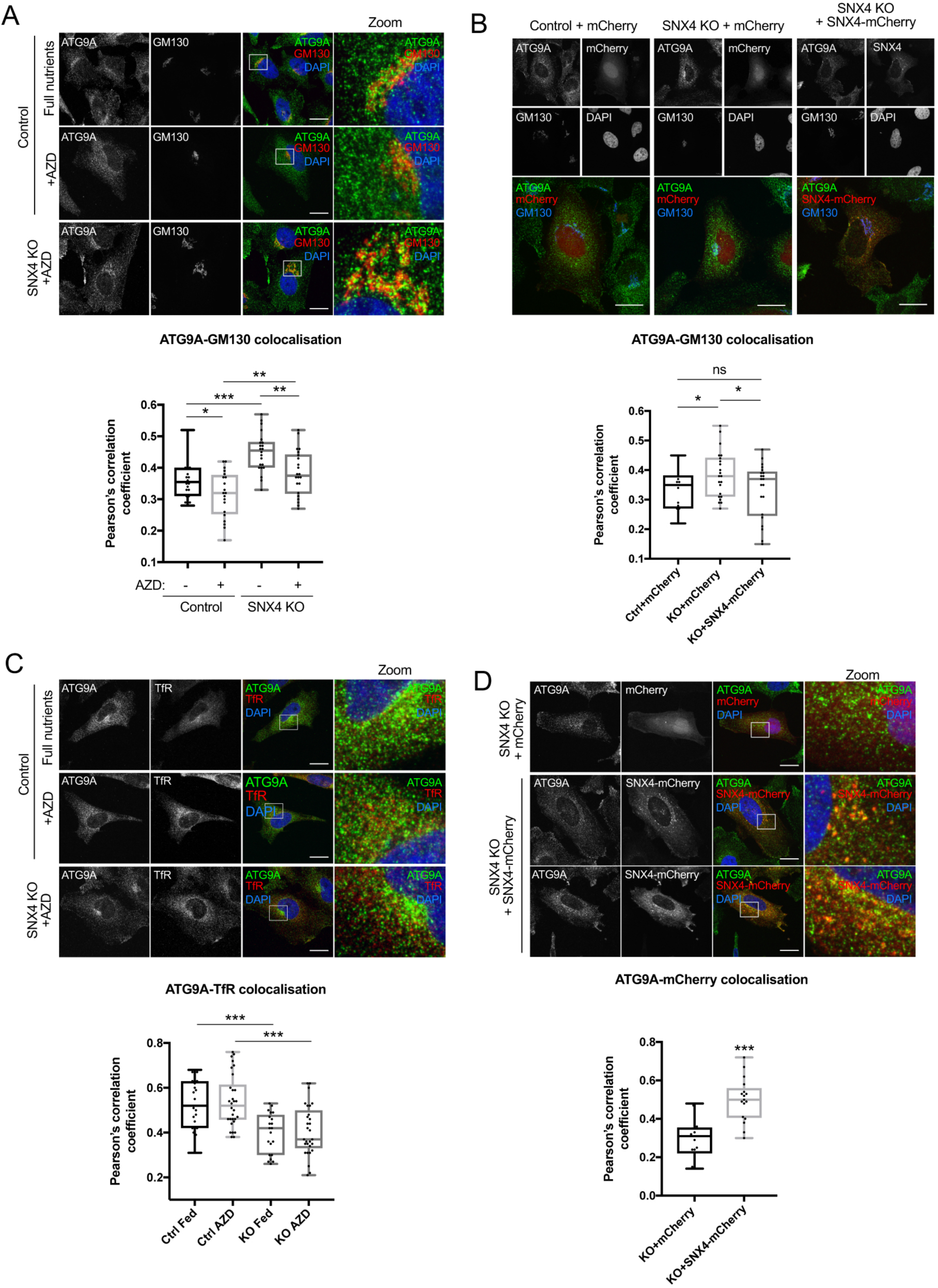
Altered ATG9A trafficking in SNX4 CRISPR KO HeLa cells. (A) ATG9A/GM130 colocalisation. Example images above; Pearson’s correlation below in cells treated or not with AZD8055. (B) Rescue of the ATG9A Golgi localisation defect in AZD8055-treated SNX4 CRISPR KO cells using mCherry-SNX4. Example images above; Pearson’s correlation below. (C) ATG9A/TfR colocalisation. Example images above; Pearson’s correlation below in cells treated or not with AZD8055. (D) mCherry-SNX4/ATG9A colocalisation in AZD8055-treated SNX4 CRISPR KO cells. Example images above; Pearson’s correlation below. *p<0.05; **p<0.01; ***p<0.001. Bars = 10 µm.

## DISCUSSION

Here, we have used an array of experimental protocols to describe how SNX4 acts as the core component of two heterodimeric complexes—SNX4:SNX7 and SNX4:SNX30—that are associated with overlapping early endosomal compartments. These findings extend the concept of specific BAR domain-mediated associations within this family of proteins, complementing the known homodimeric interactions observed within the SH3 domain-containing SNX-BARs, SNX9, SNX18 and SNX33 (Dislich et al., 2011; Haberg et al., 2008; van Weering et al., 2012); but also see (Park et al., 2010)), and the heterodimeric assemblies that comprise the SNX-BAR membrane deforming ESCPE-1 coat complex, SNX1, SNX2, SNX5, SNX6 (and SNX32) (Rojas et al., 2007; Simonetti et al., 2019; Wassmer et al., 2007; Wassmer et al., 2009). The molecular interactions that govern BAR domain homo-versus heterodimerisation remain opaque. Molecular modelling of SNX33 homodimers (Dislich et al., 2011), based on the published X-ray crystallographic structure of the SNX9 homodimer (Pylypenko et al., 2007), is consistent with the presence of a large dimer interface that contains a number of salt bridges, hydrophobic interactions, and hydrogen-bonding networks that together support tight homodimer formation (Dislich et al., 2011; Pylypenko et al., 2007). The lack of conservation between SNX9 and SNX33 in many of the amino acids that generate these interaction networks would appear sufficient to favour homo-over heterodimerisation in this context (Dislich et al., 2011). Consistent with this, molecular modelling and an X-ray crystallographic structure of the SNX1 homodimer has established how SNX1:SNX5 heterodimer formation arises (van Weering et al., 2012). One would predict, therefore, that the organisation of a similar network of interactions would favour the formation of the described SNX4 complexes, whilst disfavouring SNX7 and SNX30 homo- and heterodimers. Unfortunately, with the lack of sequence homology between the BAR domain of SNX1 and SNX9 and the corresponding domains in SNX4, SNX7 and SNX30, atomic resolution structures of the various SNX4 complexes will be required to describe how this dimer interface is organised.

In mammalian cells, SNX4 has been implicated in the tubular-based sorting of internalised TfnR from early endosome to the endocytic recycling compartment (Traer et al., 2007), and in the endosome-to-Golgi transport of ricin (Skanland et al., 2009; Skanland et al., 2007). Our data reveal that a portion of SNX4 colocalises with ATG9A in the Golgi region, and coordinates its trafficking to establish productive autophagosome assembly sites in mammalian cells. In yeast, Snx4 associates with the PAS (Zhao et al., 2016), via binding to Atg17 (Nice et al., 2002; Uetz et al., 2000), and supports efficient selective (see (Lynch-Day and Klionsky, 2010)) and non-selective (Ma et al., 2018) forms of autophagy. We observed partial and transient correlations between SNX4 and markers of mammalian autophagosome assembly sites (**Fig. 1D; Videos 1 & 2**). Despite this, we did not find evidence for SNX4 association with the mammalian equivalent of Atg17 (FIP200; or indeed any other core autophagy protein), either by pull-downs (**Fig. S6**) or by quantitative SILAC proteomics (data not shown), suggesting that its influence on mammalian autophagy may be indirect. To determine where the defect in non-selective autophagy occurs in cells deficient for SNX4, we first carried out a series of imaging-based experiments in fixed and live cells siRNA silenced for *SNX4, SNX7* and/or *SNX30*, leading to 2 important observations: (i) that SNX4:SNX7 is the mammalian autophagy SNX4 heterodimer (e.g. **Fig. 2C, Fig. 4F-I, Fig. 5C**); and that the autophagy deficiency in SNX4 and/or SNX7 suppressed cells arises at the point of ATG5 recruitment, upstream of ATG8 (LC3) lipidation (e.g. **Fig. 4F-I, Fig. 5C-F**). In yeast, Snx4 acts with two additional SNX-BARs, Snx41/Atg24 and Snx42/Atg20, to mediate retrieval of the SNARE Snc1 from post-Golgi endosomes back to the late Golgi (Hettema et al., 2003). Given that yeast Snx42/Atg20 is the expected SNX30 equivalent (Popelka et al., 2017), and with evidence that a Snx4:Atg20 dimer is needed for lipid homeostasis to support efficient autophagosome-to-vacuolar fusion (Ma et al., 2018), it is possible that yeast and mammalian cells differ in their mechanistic requirements for either SNX30/Snx42 or SNX7/Snx41 acting in concert with SNX4/Snx4 during autophagy. Indeed, evidence suggests that the yeast Snx4:Snx42 complex may act at later stages of autophagy (i.e. at the vacuolar fusion step (Ma et al., 2018)), and/or specifically during selective autophagy, meaning that further studies focused on the possible roles of SNX30/Snx41 are merited.

The molecular basis for the influence of the SNX4:SNX7 heterodimer during early stages of mammalian autophagy remains to be fully elucidated. Our data point strongly to a requirement for SNX4:SNX7 at the ATG12∼ATG5 recruitment stage, because GFP-ATG5 puncta are relatively short lived and generally less bright in cells lacking SNX4 when compared with control cells (**Fig. 5D-F**); meanwhile in SNX7 depleted cells, autophagosome assembly site markers including ATG13, ATG5, WIPI2 and ATG16L1 are clearly and dramatically amplified, arguing for a stalling of the autophagosome assembly pathway (**Fig. 4E-I**). Despite this, LC3B lipidation and/or puncta formation was only partially dampened following SNX7 suppression (e.g. **Fig. 4B**), perhaps indicating that increasing the abundance of less efficient assembly sites balances the suppressed autophagy response at the level of LC3B lipidation. Given that autophagic flux in SNX7/SNX30 suppressed cells (as measured by p62/SQSTM1 turnover kinetics; **Fig. 4C**) appeared to be somewhat affected, it remains possible that these SNX-BARs have partially redundant autophagy roles in partnership with SNX4, at both assembly and maturation stages. This would be consistent with our observation that ATG5 puncta number/kinetics differ when comparing SNX4 and SNX7 siRNA suppressed cells which would not be expected should they be acting wholly in concert and as the sole SNX4-containing mammalian autophagy heterodimer (**Fig. 4H, 5D-F**); however, different siRNA silencing efficiencies and the likely impact of an altered SNX4:SNX7/SNX4:SNX30 heterodimer balance when siRNA suppressing either partner SNX-BAR cannot be excluded.

The Simonsen lab has shown that another SNX-BAR, SNX18, acts as a positive regulator of mammalian autophagy (Knaevelsrud et al., 2013; Soreng et al., 2018). SNX18 is targeted to the recycling endosome from where it directs ATG16L1 and LC3-positive membrane delivery to the autophagosome assembly site (Knaevelsrud et al., 2013). This role requires the membrane tubulation capabilities of SNX18, and is promoted by an LC3-interacting (LIR) motif identified in the SH3 region of SNX18 (Knaevelsrud et al., 2013). In further work, SNX18 was found to regulate autophagy via mobilisation of ATG9A-positive vesicles from the recycling endosome, with a requirement for SNX18 binding to dynamin-2 (Soreng et al., 2018). Clearly there are similarities between our characterisation of SNX4:SNX7-mediated autophagy and the involvement of SNX18 in the same pathway, although neither SNX4 nor SNX18 were identified as autophagy regulators in the respective, alternate siRNA-based screens (this study; (Knaevelsrud et al., 2013)), perhaps due to cell-line differences and/or siRNA efficiencies. At the level of autophagosome assembly kinetics, important differences were observed between these autophagy-regulating SNX-BARs. For example, whilst siRNA depletion of either SNX-BAR reduced the autophagy response in both studies, LC3 lipidation was enhanced following SNX18 overexpression (Knaevelsrud et al., 2013), but was suppressed when SNX4 was overexpressed (**Fig. 1F**). Surprisingly, there were no clear differences in LC3 lipidation levels in SNX18 KO cells except when assessed in full media in the presence of BafA1, although long-lived protein turnover was significantly reduced (Soreng et al., 2018). This differs from the LC3 lipidation defects observed in acute SNX18 siRNA-depleted cells (Knaevelsrud et al., 2013), suggesting that a form of functional compensation may have arisen. Tellingly, this is also distinct from the scenario we describe in SNX4 siRNA depleted and CRISPR KO cells where LC3B lipidation is robustly suppressed in both cases (this study). Elsewhere, in starved SNX18 KO cells, WIPI2 and ATG16L1 puncta numbers were significantly lower (Soreng et al., 2018), whereas SNX4 and/or SNX7 siRNA suppression dramatically elevated WIPI2 and ATG16L1 puncta numbers during starvation (this study). Further important differences were revealed upon analysis of ATG9A dynamics within SNX18 and SNX4 KO cells: whereas SNX18 KO increased colocalization between ATG9A and TfR (Soreng et al., 2018), this was significantly reduced in SNX4 KO cells (**Fig. 7C**). Indeed, in SNX4 KO cells, ATG9A was more strongly associated with the Golgi region under basal conditions and during autophagy induction (**Fig. 7A**). Interestingly, colocalization between ATG16L1 and WIPI2 was observed to be weakened by SNX18 KO (although not statistically significant; (Soreng et al., 2018)), a situation also seen in SNX7 siRNA depleted cells at the level of WIPI2/ATG5 colocalisation (**Fig. 5A, C**). Evidently, these autophagy-regulating SNX-BARs have common influences at the ATG16L1 and/or ATG5 recruitment stage; a finding that is consistent with the autophagy defects observed in ATG9A KO cells (e.g. (Orsi et al., 2012)).

## MATERIALS AND METHODS

### Materials and antibodies

All materials were purchased from Sigma unless otherwise stated. BafA1 (B1793); AZD8055 (Seleckchem, S1555); CHX (C7698); puromycin (P8833); DAPI (D121490). The following antibodies were used: anti-GAPDH (G8796); anti-ATG9A (Abcam, ab108338); anti-ATG16L1 (MBL, PM040); anti-LC3B (L7543); anti-ULK1 (Cell Signalling, D8H5/8054); anti-ATG7 (Cell Signalling, D12B11/8558); anti-ATG3 (Cell Signalling, 3415); anti-FIP200 (SAB4200135); anti-GFP (Covance, MMS-118P; 1:2000 IB); anti-GM130 (Santa Cruz, sc216); anti-p62/SQSTM1 (Abnova, H00008878); anti-WIPI2 (BioRad, MCA5780GA); anti-SNX4 (Abcam, ab198504); anti-SNX30 (Abcam, ab121600); anti-SNX7 (Proteintech, 12269-1-AP); anti-SNX1 (BD Biosciences, 611482); anti-SNX2 (BD Biosciences, 611308); anti-SNX4 (Santa Cruz, sc-10623); anti-SNX5 (Proteintech, 17918-AP); anti-SNX6 (Santa Cruz, sc-8679); anti-SNX7 (Abcam, Ab37691); anti-EEA1 (BD Biosciences, 610456); anti-CD63 (Santa Cruz, sc-51662); anti-APPL1 (kind gifts from Pietro De Camilli, Yale University and Philip Woodman, University of Manchester); anti-β-actin (A1978); anti-TfR (Santa Cruz, sc-65882); anti-mouse HRP (Stratech, G32-62DC-SGC); anti-rabbit HRP (Stratech, G33-62G-SGC); Alexa Fluor 488 (Invitrogen, A-11029/A-11034); Alexa Fluor 568 (Invitrogen, A-11031/A11036); Alexa Fluor 647 (Invitrogen, A-21236/A-21244).

### Cell-lines and cell culture

Parental HeLa and hTERT-RPE1 cells, GFP-ATG13 RPE1 (this study), ATG5-GFP RPE1 (MacVicar et al., 2015), YFP-LC3B RPE1 (this study), SNX4-GFP HeLa (this study), SNX4-GFP hTERT-RPE1 cells (this study), GFP-LC3B HEK293 (a gift from Sharon Tooze) stable cell-lines, and the SNX4 CRISPR null HeLa cells (including wild type and CRISPR KO cells stably expressing mCherry-GFP-LC3B; this study) were maintained in DMEM containing 4500 mg/ml glucose (Sigma, D5796), supplemented with 10% FBS (Sigma, F7524); (with 1% penicillin-streptomycin [Sigma, P4333] for CRISPR null cell-line cloning and maintenance, only). Cells were grown at 37°C with 5% CO_2_.

### Vector design and cloning of SNX-BARs, viruses and transductions

The SNX-BAR genes were amplified from a HeLa cell cDNA library using conventional PCR. For transient transfection of mammalian cells, the SNX-BAR genes where cloned into pEGFP.C1 and pmCherry.C1 vectors, for the transduction of mammalian cells the genes where cloned into EGFP/mCherry.C1 pXLG3 lentivector system. Lentiviruses were generated in HEK293T cells by transfection with cDNAs along with packaging vectors pMD2G and pAX2. Lentiviral particles were collected at 48 h, cleared by centrifugation (2900 × *g*; 10 min), then passed through a 0.45 µm polyethersulfone filter to be used immediately or stored at −80°C. Control or SNX4 suppressed HeLa cells overexpressing mCherry or mCherry-SNX4 were generated as follows: cells were plated in 6-well plates and transduced with the corresponding lentiviruses. After three days, cells were seeded on coverslips for the corresponding experiment.

### CRISPR/Cas9 for generation of SNX4 KO cells

The CRISPR-Cas9 plasmid developed by the Zhang lab (pX330) was used for targeted gene knockout (Cong *et al*., 2013). The sequences for the gRNAs (5’-GCGGTCGGCAAGGAAGCGGA-3’) were calculated using the online tool from the Zhang Lab and cloned in pX330 accordingly (www.genome-engineering.org). To generate SNX4 knock-out cells, Hela cells were co-transfected with CRISPR-Cas9 plasmids and a plasmid conferring puromycin resistance using FuGENE (Promega, E2693). 24 h after transfection, cells were subjected to 24 h of puromycin selection (Sigma, P8833; 2 µg/ml), after which cells were resuspended using accutase (BioLegend, 423201) and diluted in Iscove’s Modified Dulbecco’s Medium (Sigma, I3390) supplemented with 10% FBS to 3.5 cells/ml. Subsequently, 200 μL cell suspension was plated in each well of ten 96-well plates, and after three weeks the plates were screened for the presence of cell colonies. Colonies were expanded in DMEM and subjected to lysis and western blotting to determine the expression levels of the target protein.

### Immunoblotting

Cells grown on 6-well plates were initially washed with ice-cold PBS, then lysed with 200 µl/well ice-cold radioimmunoprecipitation assay (RIPA) buffer consisting of: 50 mM Tris HCl (pH 7.4); 1% Triton-X-100 (Sigma, 9002-93-1); 0.5% sodium deoxycholate (Sigma, D6750); 150 mM NaCl (Sigma, S9888); 0.1% sodium dodecyl sulphate (SDS) (Sigma, 436143); supplemented with one tablet of protease inhibitor per 10 ml of RIPA buffer. The homogenates were incubated on ice for 10 min, then cleared by centrifugation at 12,000 × *g* for 15 min at 4°C. Supernatants were collected as soluble fractions. Proteins were transferred to nitrocellulose membranes (BioRad, 1620115), and membranes were then incubated overnight with primary antibody diluted in 5% milk or 2.5% BSA in Triton X-100-TBS buffer. Primary and secondary antibodies used are listed above. Membranes were then washed three times prior to incubation with ECL Chemiluminescence reagents (Geneflow, K1-0170), and band intensities were detected on film (GE Healthcare, 28906837).

### GFP-Trap

For GFP-Trap immunoisolation, cells in 10 cm plates were washed with ice cold PBS, and 500 µl of lysis buffer (50 mM Tris base, 0.5% NP40, 1 mM PMSF, 200 µM Na_3_VO_4_, protease inhibitors, pH 7.5) was added. Cells were scraped and lysates collected and incubated on ice for 10 min. Lysates were cleared by centrifugation at 20000 × *g* for 10 min at 4°C and added to pre-equilibrated beads (Chromotek, GTA-100) and rotated for 2 h at 4°C. The sample was then spun at 2700 × *g* for 2 min at 4°C to pellet beads, which were washed 3x in wash buffer (50 mM Tris base; 1 mM PMSF; 200 µM Na_3_VO_4_; protease inhibitors) then resuspended in SDS-PAGE gel sample buffer.

### Immunoprecipitations

Confluent HEK293 cells in 15 cm dishes were washed with ice-cold PBS at least three times prior to lysis. Cells were lysed with 500 µl of ice-cold lysis buffer (50 mM Tris-HCl, 0.5% NP-40, and Roche protease inhibitor cocktail, pH7.5) and lysis was aided through the use of a cell scraper. Cell lysates were cleared by centrifugation in a bench-top centrifuge for 10 min at 13,000 rpm at 4°C. Cleared lysates were incubated with 2 µg of either IgG control antibody or SNX1, SNX4, SNX7 and SNX30 antibody overnight at 4°C on a roller. 25 µl of cleared lysate was retained as 5% total protein input. Protein G sepharose beads (GE Healthcare) were washed three times in lysis buffer to remove any residual ethanol from the storage buffer. Pre-equilibrated Protein G sepharose was then added to the Eppendorf tubes containing lysate/antibody mixture and these tubes were further incubated for 1 h at 4°C on a roller. After incubation, the beads were pelleted by centrifugation at 4 °C for 30 seconds at 4000 rpm. The beads were then washed 3 times in 1 ml of lysis buffer. Finally, all buffer was removed and Protein G beads (and associated immuno-precipitated proteins) were either stored at −20°C or processed immediately for SDS-PAGE and Western blot analysis.

### siRNA suppression

The following siRNA oligonucleotides were used for experiments targeting SNX4/SNX7/SNX30 proteins in autophagy (Dharmacon, siGENOME or SMARTpool): NT#1 (gacaagaaccagaacgcca); NT#2 (guacgcggaauacuucgauu); ATG5 (ggaauauccugcagaagaa); SNX4 SMARTpool D1 (uuacugaccuuaagcacua), D2 (gaaacaaggucaguugaac), D3 (gcggcgauauagugaauuu), D4 (gcgacggauugguuuagaa); SNX7 SMARTpool D1 (gcggaugucuggacucuca), D2 (guacgugcuuuauagugaa), D3 (ggagacgauaucaagauuu), D4 (gcacaccccacucugauua); SNX30 SMARTpool D1 (acaagaacauccaguauua), D2 (gaagaagagggaccaaguu), D3 (cggcggacgucgagaaaug), D4 (ggagucgauuauuccacua).

### Immunofluorescence and cell imaging

For fixed cell imaging, cells were seeded on coverslips, washed twice with PBS and incubated with 4% formaldehyde for 15 min or −20°C methanol for 5 min. Cells were then incubated for 30 min with primary antibodies (listed above) in PBS. Cells were washed three times with PBS and incubated with secondary antibodies (listed above) and counterstained with DAPI (100 ng/ml) for 10 min. Cells were then washed again with PBS and mounted in Mowiol containing 25mg/ml DABCO (1,4-Diazabicyclo [2.2.2] octane; D27802). Fixed-cell images were obtained using an Olympus IX-71 inverted microscope (60x Uplan Fluorite objective; 0.65-1.25 NA, oil immersion lens) fitted with a CoolSNAP HQ CCD camera (Photometrics, AZ) driven by MetaMorph software (Molecular Devices). Confocal microscopy was carried out using a Leica SP5-AOBS confocal laser scanning microscope (63x oil immersion objective, 1.4 NA; or 100x oil immersion objective, 1.4 NA) attached to a Leica DM I6000 inverted epifluorescence microscope. Laser lines were: 100 mW Argon (for 458, 488, 514 nm excitation); 2 mW Orange HeNe (594 nm); and 50 mW diode laser (405 nm). The microscope was run using Leica LAS AF software (Leica, Germany). MetaMorph software was used to quantify puncta numbers. A TopHat morphology filter was used to score circular objects of 5 pixels (∼1 µm) diameter. An automated cell count was then performed to count the number of selected items. For a typical experiment, fifteen random fields were imaged and puncta numbers per cell in each field was counted. Colocalization was determined by acquiring images from ∼25 cells per condition, with Pearson’s coefficient calculated using Fiji software.

### Directed yeast two-hybrid screens

The yeast strain AH109 was co-transfected with bait vector full-length human SNX4 or Lamin cloned into pGBKT7 (Clontech, Oxford, UK) against a pray library of FL SNX-BARs. Yeast clones with positive bait–prey interactions were selected on SD –Leu –Trp plates supplemented with 1 mM 3′ AT and α-X-Gal (Glycosynth, Cheshire, UK).

### Statistical analysis

Graphical results were analyzed with GraphPad Prism 7 (GraftPad Software, San Diego, CA), using an unpaired Student’s t-test (where not specified), or a Kruskal-Wallis test for non-parametric ANOVA. Results are expressed as mean ± SEM or mean ± SD, as indicated.

## AUTHOR CONTRIBUTIONS

The project was conceived and designed by JDL and PJC. Initial autophagy SNX screens were carried out by NA. VMSB carried out all of the autophagy-related siRNA and overexpression work comprising Figs 1, 4, 5, S1, S2, S3, S6A,B. CJT performed the yeast 2-hybrid, SNX IP and SNX siRNA analysis, as well as the SNX endosomal imaging comprising Figs 2 & 3. BS contributed Fig. S4, designed the SNX4 CRISPR guide RNAs, and carried out CRISPR pilot experiments; ZA generated the SNX4 CRISPR KO cells and characterized the autophagy response in these cells, contributing Figs 6, 7, S5, S6C. JDL and PC analysed data, and JDL wrote the manuscript.

## ACKNOWLEDGEMENTS

This work was funded by the BBSRC (BB/J002704/1), the BBSRC/EPSRC through the BrisSynBio Synthetic Biology Research Centre (BB/L01386×1), and the Wellcome Trust (089928/Z/09/Z). We are grateful for the support of the Wolfson Bioimaging Facility at Bristol University.

