## Supplemental figs for "A heterodimeric SNX4:SNX7 SNX-BAR autophagy complex coordinates ATG9A trafficking for efficient autophagosome assembly"

**SUMMARY STATEMENT:** A heterodimeric SNX4:SNX7 SNX-BAR complex regulates mammalian autophagosome assembly through the control of ATG9 trafficking.

### SUPPLEMENTAL FIGURES

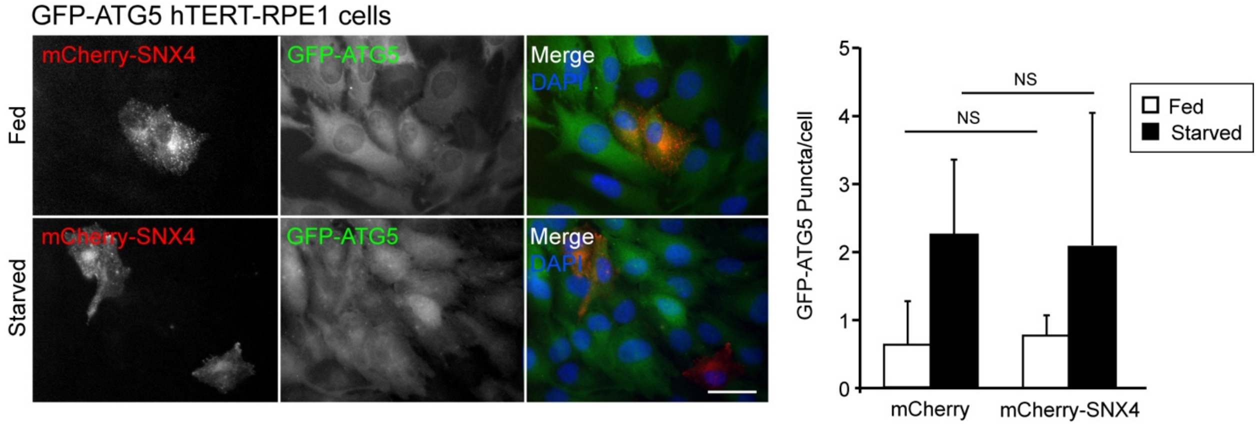


**Figure S1:** Steady state ATG5 puncta numbers are not altered in cells overexpressing SNX4. ATG5-GFP hTERT RPE1 cells were transiently transfected with mCherry-SNX4 (mCherry as control), and ATG5 puncta numbers were counted using automated software (MetaMorph) in fed and starvation conditions (1 h). Example images of mCherry-SNX4 fields to the left; quantitation to the right. Mean ± SD; n=3; Bar = 20 μm.

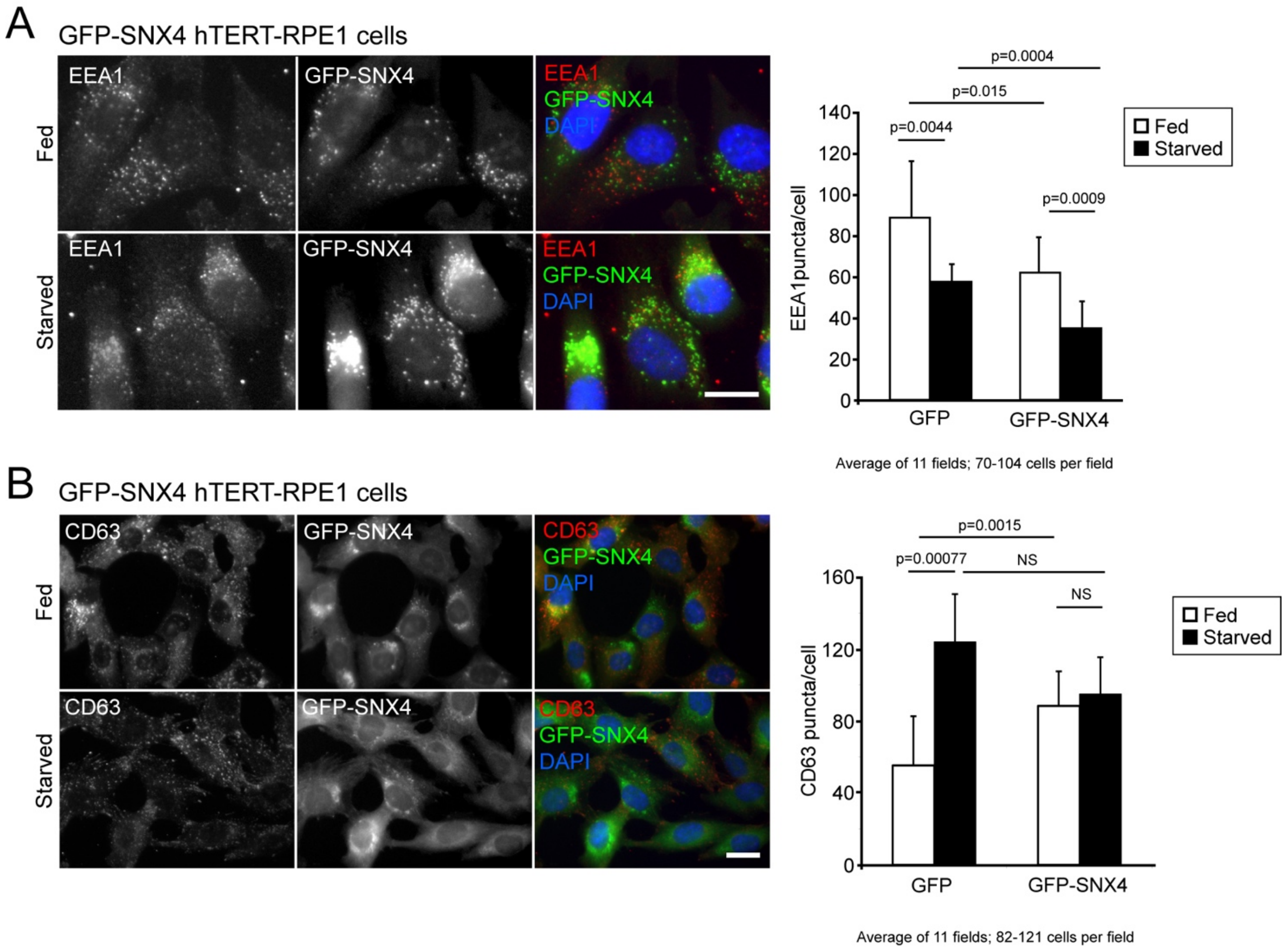


**Figure S2:** GFP-SNX4 stable hTERT RPE1 cells show defects in endolysosomal compartments. (A) hTERT RPE1 cells stably overexpressing GFP-SNX4 were starved (1 h), then fixed and stained with antibodies against the (**A**) early endosome (EEA1) and (**B**) the lysosome (CD63). Example images to the left; quantitation to the right. Means ± SD; p values are shown on the graphs. Bars = 10 μm.

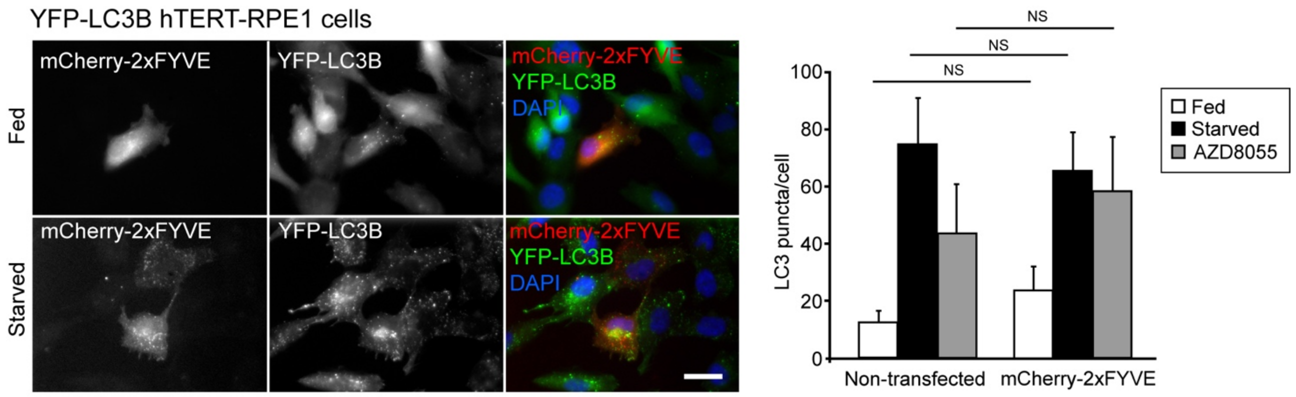


**Figure S3:** Overexpression of 2xFYVE does not influence the autophagy response. hTERT RPE1 cells stably expressing YFP-LC3B were transiently transfected with mCherry-2xFYVE, and LC3B punta were counted following starvation or AZD8055 treatment (1 h) in transfected and untransfected cells. Example images of fed and starved cells to the left; quantitation to the right. Bar = 10 µm.

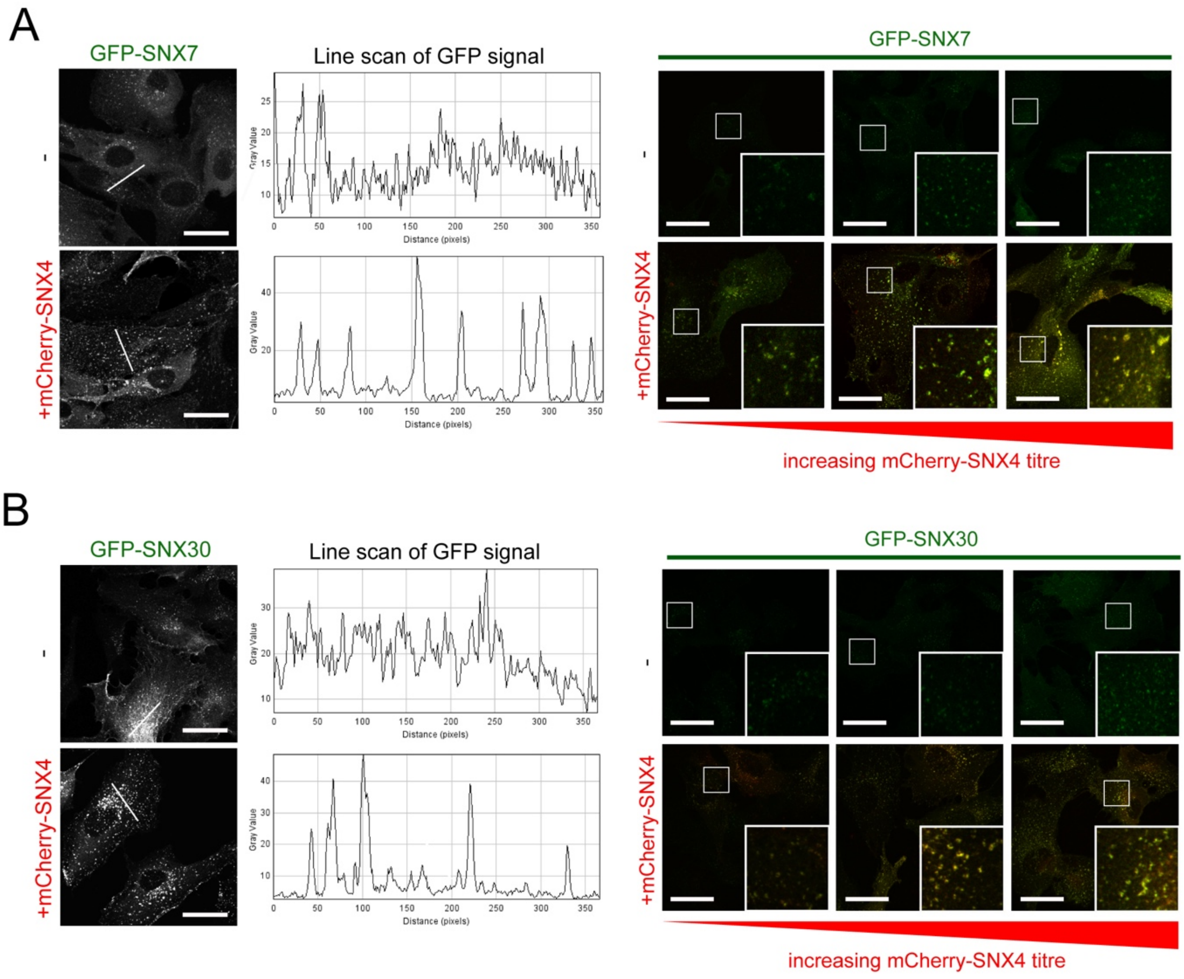


**Figure S4:** GFP-SNX7 and GFP-SNX30 display predominant cytoplasmic localisations when expressed alone; however, when co-expressed with mCherry-SNX4 they show a punctate localisation that correlates with SNX4 levels. RPE1 cells were lentivirally transduced with GFP-SNX7 (**A**) or GFP-SNX30 (**B**), alone or in combination with increasing titres of mCherry-SNX4. Transduced cells were then fixed and imaged. Line scans of GFP signal show that GFP-SNX7 and GFP-SNX30 localise on cytosolic puncta only when the heterodimeric partner SNX4 is co-expressed. Bar = 20 μm.

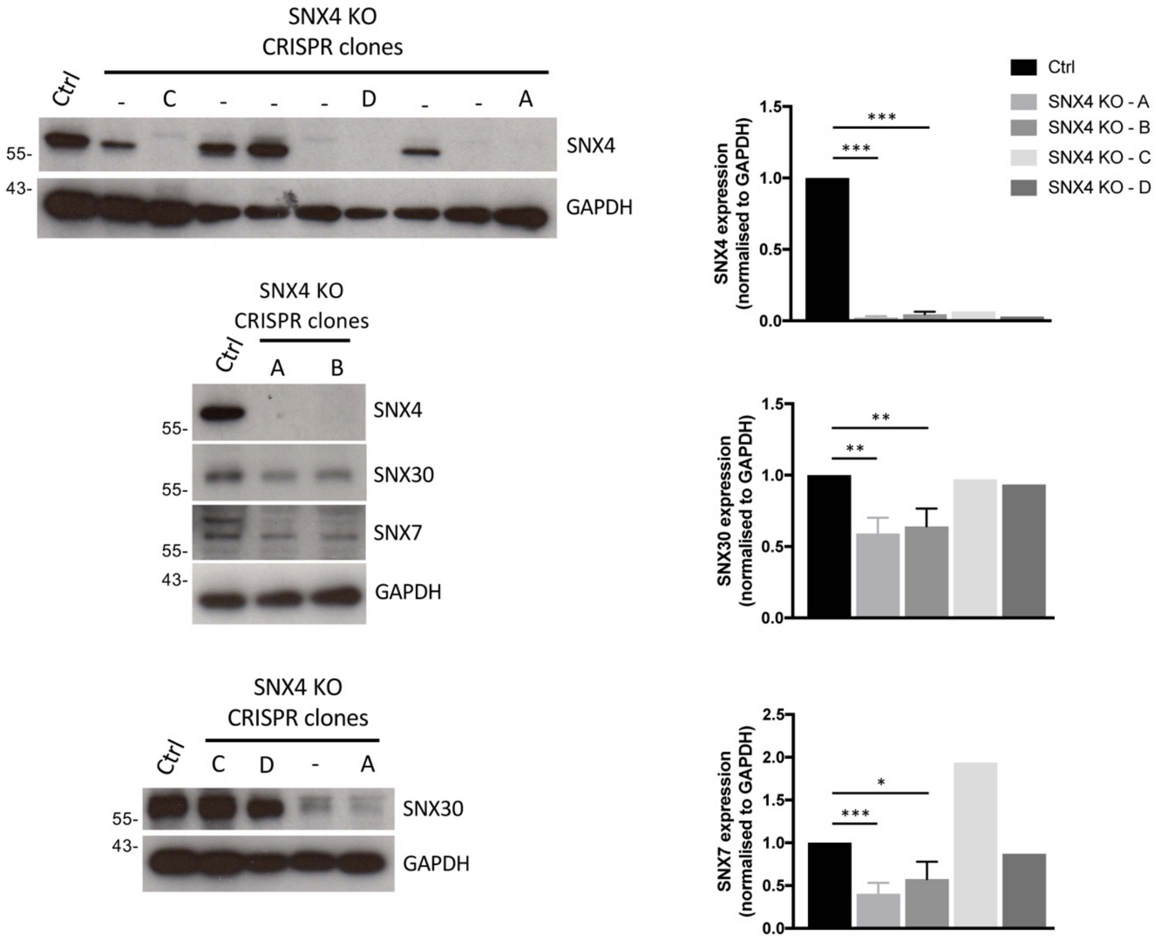


**Figure S5:** Analysis of SNX4 CRISPR clones. Individual colonies of HeLa cells following SNX4 CRISPR-Cas9 treatment were selected and expanded. Example blots for SNX4, SNX7, and SNX30 are shown to the left, with quantitation of clones A-D shown to the right. *p<0.05; **p<0.01; ***p<0.001.

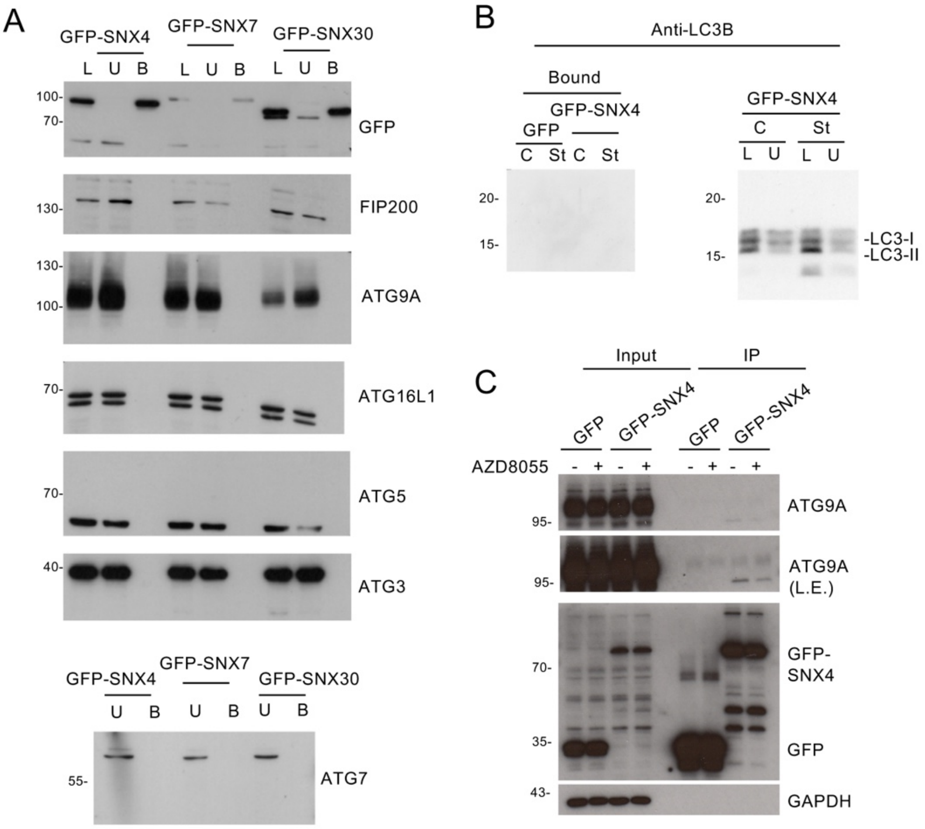


**Figure S6:** SNX4/7/30 do not co-precipitate with known autophagy proteins. (**A**) HeLa cells expressing GFP-SNX4/7/30 were lysed and subjected to GFP-TRAP immunoprecipitation. Lysates (L), unbound fractions (U) and bound fractions (B) were blotted for the autophagy markers shown. (**B**) GFP-SNX4 HeLa cells were starved (1 h), fractionated, and lysates used for GFP-TRAP immunoprecipitation. Samples were blotted for LC3B, which was not detected in the bound fractions for GFP-SNX4 or GFP (control) lysates. (**C**) GFP-SNX4 cells were treated with AZD8055, lysed, and subjected for GFP-TRAP immunoprecipitation. Inputs and bound fractions were blotted for ATG9A. (L.E. = long exposure).
